# Different approaches to estimate benthic metazoan diversity associated with free-living macroalgae (*Fucus vesiculosus*) on shallow soft sediments

**DOI:** 10.1101/2023.02.07.527458

**Authors:** Roxana Preston, Markus Majaneva, Viivi Halonen, Iván F Rodil

## Abstract

Habitat complexity can boost biodiversity by providing a wide range of niches allowing species co-existence. Baltic Sea benthic communities are characterised by low species diversity. Thus the occurrence of the habitat forming macroalga *Fucus vesiculosus* may influence benthic communities and promote diversity. Here we obtain biodiversity estimates through conventional and eDNA approaches for the benthic assemblages associated with free-living *Fucus* and the adjacent bare-sediment habitats at six sites from the Northern Baltic Proper and the Gulf of Finland. Free-living *F. vesiculosus* habitats are heterogeneous with biodiversity estimates varying considerably among sites. The additional habitat complexity provided by *F. vesiculosus* tends to improve taxa richness as a result of additional epifauna assemblages, although infaunal taxa richness and abundance is often reduced. Consequently the complex habitats provided by free-living *F. vesiculosus* often improves biodiversity, yet alters the composition of assemblages in soft sediment habitats and consequential ecosystem functioning. We emphasise the disparity in biodiversity estimates achieved when employing different biodiversity approaches. Biodiversity estimates were more similar within approaches compared to between habitat types, with each approach detecting exclusive taxa. We suggest that biodiversity estimates benefit from a multi-approach design where both conventional and eDNA approaches are employed in complement.

## 1 Introduction

Habitat complexity influences biodiversity and associated ecosystem functioning (Kovalenko et al. 2012). In many ecosystems higher habitat complexity will attract more associated species (Johnson and Agrawal 2005) by promoting species coexistence through providing a wide range of niches, thereby reducing niche overlap and increasing diversity (Huston and DeAngelis 1994; Levins 1979). The macroalgae genus *Fucus* represents important foundation species within northern hemisphere coastal environments. Within the Baltic Sea, *Fucus vesiculosus* (herein *Fucus*), is one of only a few large, perennial macroalgae forming structurally complex canopies in the coastal photic zone, supporting numerous associated organisms (Henseler et al. 2019; Kraufvelin and Salovius 2004; Wikström and Kautsky 2007). Attached *Fucus* canopies are some of the most highly productive habitats within the Baltic Sea (Attard et al. 2019).

Alongside the typical attached form, a free-living form is common throughout the Baltic Sea on any substrate type, although most frequently on soft sediments, in more sheltered areas (Preston et al. 2022a, 2022b). Consequently the free-living form can form stable, perennial mats in locations where attached algae would otherwise be absent. These free-living mats create three-dimensional habitats of varying heights and densities that are comparable to the interstitial space of sediments (HELCOM 2013). Thus free-living *Fucus* likely provide high complexity habitats alongside also influencing the normally associated fauna of soft sediment habitats. In fact algal mats are often associated with high biodiversity (El-Khaled et al. 2022; Rossbach et al. 2022, 2021). As Baltic Sea benthic habitats are characterised by exceptionally low species diversity (Kotta and Orav 2001) the addition of algal cover may provide several conditions, including increasing food resources (Arroyo et al. 2006; Norkko et al. 2000) and providing protection from predation (Aarnio and Mattila 2000; Norkko et al. 2000), which may in turn boost biodiversity. Although algal cover may also contribute to hypoxic conditions within the sediment resulting in faunal reductions (Everett 1994; Norkko and Bonsdorff 1996a, 1996b; Rabalais et al. 2010).

Baseline biodiversity estimates for benthic habitats can be obtained through conventional approaches whereby samples are collected by sampling devices (e.g. cores, quadrats, scuba diving), sorted and individually taxonomically identified. This approach is often time-consuming and requires specialist taxonomic expertise. Increasingly, DNA-based molecular techniques are being utilised to assess biodiversity (Zaiko et al. 2018). Environmental DNA (eDNA) originates from living organisms, dead cells and extracellular DNA present within the sample (Levy-Booth et al. 2007; Pietramellara et al. 2009; Taberlet et al. 2012). DNA sequence information from the pool of genetic material within the environmental sample is used to determine the taxonomic identification within the sample. Here we utilise these two approaches to provide biodiversity estimates for two habitat types: *Fucus* associated soft sediments and the adjacent bare-sediment. Firstly we apply two conventional sampling approaches to identify macroinfaunal and epifaunal benthic communities and secondly an eDNA approach on sediment samples to identify metazoan benthic assemblages. The aims of the study are to (i) investigate the influence of free-living *Fucus* on soft sediment faunal benthic assemblages, including the spatial variability among sites and countries, and (ii) evaluate the discrepancies between approaches to biodiversity estimates.

## 2 Methods

### 2.1 Sample locations

The Baltic Sea is a semi-enclosed brackish system with a defined north-south salinity gradient (Furman et al. 2014; Lüning 1990; Zillén et al. 2008). Surface salinity ranges from 8–10 in the southern Baltic, 7–8 in the Baltic Proper and down to 3–6 in the Gulfs of Bothnian and Finland (Lüning 1990; Matthäus 2006). Species diversity is low, having approximately 10 times fewer species compared to the neighbouring North Sea (Elmgren and Hill 1997; Johannesson et al. 2011). Sampling was performed at two locations approx. 340 km apart: Askö in the Northern Baltic Proper and Tvärminne in the Gulf of Finland (Figure 1). Three sites were selected per location (Table 1). All sites were within close proximity of the shore, in shallow, sheltered embayments associated with *Phragmites australis* reed beds. The bottoms at all sites were soft, being either clay, sandy or mixed substrata. Free-living *Fucus* was the dominant macroalga within these locations. At sites AS1 and TZ1 the thalli were entangled within *P. australis*. Salinity at the sites ranged from c. 5.9–6.0 at Askö and c. 5.9–6.1 at Tvärminne whilst maximum depth ranged from 1.9–3.4 m at Askö and 2.5–3.2 m at Tvärminne.

**Figure 1:**
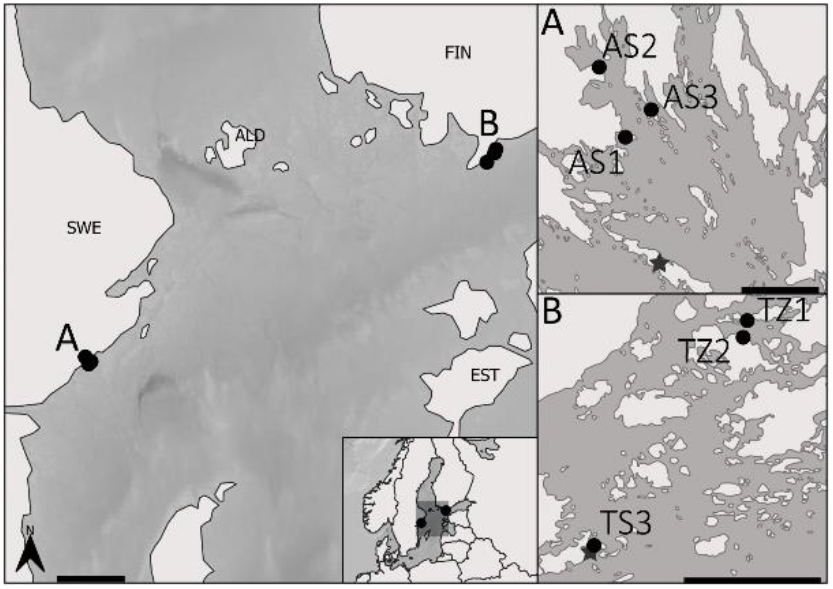
Locations of the sampling sites. A: Askö, B: Tvärminne. Scale bars represent 50 km in the main map and 5 km in the inset maps (A and B). Star symbols represents field stations.

**Table 1:** Sampling site information

| Site | Subbasin | Country | Location | Coordinates<br>(decimal<br>degrees) | Date | Salinity | Depth<br>(m) |
| --- | --- | --- | --- | --- | --- | --- | --- |
| AS1 | Northern Baltic Proper | Sweden | Askö | 58.89455<br>17.62786 | 13/06/19 | 6.1 | 2.5 |
| AS2 | Northern Baltic Proper | Sweden | Askö | 58.93688<br>17.60736 | 13/06/19 | 6.1 | 3.2 |
| AS3 | Northern Baltic Proper | Sweden | Askö | 58.90913<br>17.66022 | 13/06/19 | 5.9 | 3.2 |
| TZ1 | Gulf of Finland | Finland | Tvärminne | 59.90994<br>23.38147 | 01/06/19 | 6 | 1.9 |
| TZ2 | Gulf of Finland | Finland | Tvärminne | 59.90469<br>23.37602 | 01/06/19 | 6 | 2 |
| TZ3 | Gulf of Finland | Finland | Tvärminne | 59.84587<br>23.25200 | 02/06/19 | 5.9 | 3.4 |

### 2:2 Sample collection and sorting

Sampling was conducted in June 2019 of two habitat types: soft sediments associated with free-living *Fucus* and the adjacent bare-sediment (Table 1). *Fucus* samples represented 100% coverage of the algae whilst bare-sediment samples lacked any form of vegetation. Samples were collected through SCUBA diving at depths ranging from 1.5–3.4 m. The conventional approach incorporated two sampling techniques to capture the macroinfauna (cores) and epifauna (quadrats) assemblages (Supplementary material S1). At each site three 20×20 cm quadrats with <1 mm mesh bags were randomly placed. Within the frame all vegetation, including epifauna, were collected. Eight benthic cores (5.6 cm diameter, 10 cm deep) were randomly collected per site (4 per habitat type). *Fucus* sediment cores were collected from underneath the free-living *Fucus* mats whilst bare-sediment cores were taken adjacent to the mats. The conventional approach used three benthic cores per habitat whilst the eDNA approach used a subsection of all four cores. The 2 ml sediment subsamples for eDNA analysis were transferred to individual microcentrifuge tubes and stored at −20 °C. The conventional macroinfaunal and epifaunal samples were run through sieves of 0.5 mm and 0.8 mm, respectively, prior to fixing in 70% ethanol. Faunal samples were sorted and identified to species level or lowest feasible taxonomic ranking. Specimens that were unable to be identified by the available taxonomic expertise were recorded as unclassified (<0.1% of total detected taxa).

### 2:3 eDNA processing & bioinformatics

Primers TAReuk454FWD1 and TAReukREV3 (Stoeck et al. 2010) targeting the 18 S nSSU gene region were used yielding fragments between 231-401 bp not including adaptors or barcodes. DNA extraction was performed using the DNeasy powersoil kit (Qiagen, 12888-100) following the standard kit protocol and stored at −20 °C. Purification of 100 μl of each DNA extract was performed using the Dneasy Powerclean Pro Cleanup Kit (Qiagen, 12997-50). DNA was quality checked on a NanoDrop™ (Thermo Scientific™) and diluted to a concentration of ~2.5–25 ng/μl. Purified samples with low yields were not diluted. Negative reactions of MQH2O and buffers from DNA extraction and purification were also processed alongside the samples. Duplicate PCR amplification was performed in 20 μl reaction mixes, with each reaction containing 10 μl Phusion Flash High-Fidelity PCR Master Mix (Thermo Scientific™, F548S), 2 μl each of forward and reverse primers (10 μM), 2 μl DNA and MQH_2_O to make up to 20 μl. Reaction were prepared on ice. The thermocycler program consisted of an initial denaturation step of 98 °C for 10 s, 10 cycles of denaturation at 98 °C for 1 s, annealing at 57 °C for 5 s and extension at 72 °C for 15 s, then 25 cycles of denaturation at 98 °C for 1 s, annealing at 47 °C for 5 s and extension at 72 °C for 15 s. A final extension of 72 °C for 2 min was performed before samples were held at 4 °C. Thermocycler programs were run on a Veriti 96-Well (Applied Biosystems). PCR products were checked by gel electrophoresis then duplicate reactions were pooled. Samples were further processed and run on Miseq (Illumina) at the DNA Sequencing and Genomics Lab, Institute of Biotechnology, Helsinki Institute of Life Science, University of Helsinki, using the 600-cycle V3 Illumina MiSeq sequencing kit.

Primers were removed from the raw amplicon reads, using cutadapt v2.1 (Martin 2011). Then, the reads were processed with the R package DADA2 1.18 (Callahan et al. 2016) in R 4.1.2 (R Core Team 2021). The quality parameters in DADA2 were adjusted based on the quality profile of the sequencing run, and were 3 maximum expected errors, 0 ambiguous bases, truncation after quality score of 13, maximum length of forward reads 211 bases, maximum length of reverse reads 201 bases, minimum overlap of 11 bases in merging and chimeric sequences were searched in consensus mode. Taxonomic affiliations of the generated ASVs (Amplicon Sequence Variants) were identified in several steps to remove unassigned and other remaining spurious ASVs. First, taxonomic affiliations were identified, using DADA2 with PR2 reference database (Guillou et al. 2013). Secondly, the PR2 database was searched, using blastn in BLAST+ 2.6.0 (Zhang et al. 2000) to identify ASVs that had low match percentage (<97%) or low query coverage (<80%) to a reference sequence in the PR2 database. Thirdly, NCBI GenBank was searched (23^rd^ May 2022), using blastn. Taxonomic affiliations of the GenBank search were parsed using the weighted lowest common ancestor algorithm in MEGAN 6.22.2 (Huson et al. 2016) with minimum bit score 600, top percentage 1.0 and minimum support 1. In the end, only ASVs that were identified to family, genus or species level were kept. Metazoan taxonomic affiliations were based on the GenBank search while all other affiliations were based on the DADA2 assignment using the PR2 database. If blastn search of the PR2 database gave low match or coverage, GenBank affiliation was used instead if family, genus or species level GenBank match was found. In addition, the ASVs that were identified to genus or family level were clustered into 97-% OTUs to represent proxies of species using vsearch v2.14.1 (Rognes et al. 2016). Finally, we removed terrestrial species of Metazoa and Embryophyta from the dataset. Four out of the eight control samples did not include any good-quality reads while three of the eight control samples included reads of species not present in the other samples (yeast *Malassezia restricta*, spider *Oecobius* sp., mite *Gamasina* sp. and an annelid worm). These species were removed from the dataset. One control sample included a single read of *Sabateria* sp. (Nematoda) that was abundant in the other samples as well, showing very low rate of tag jumps in the dataset. At site AS2 a single replicate per habitat recorded no sequences and thus these replicates were discarded. Sequence reads were normalized to relative abundance per sample.

### 2:4 Statistical tests

Counts from the conventional approach were converted into abundance (per m^2^) and then standardised as relative abundances. In the eDNA approach only metazoan taxa were included. Sequence reads were converted to relative abundances. Taxa were first standardised to phylum or class level to allow direct comparisons. All analyses were performed in R 4.0.3 (R Core Team 2021), unless expressly mentioned. Comparisons of sampling methods were made through Venn diagrams, drawn using ggVennDiagram (Gao et al. 2021). Replicates per sites were grouped and data were converted to presence/absence format. Within the Venn diagrams taxa were classified to class level. For assemblage compositions replicates per sites were grouped and plotted using ggplot2 (Wickham 2016) with taxa classified to phylum level. Multiple Response Permutation Procedure (MRPP) ordination was used to analyse the differences between *Fucus* associated assemblages across country and site using the vegan package (Oksanen et al. 2022). As direct comparison between the separate approaches were not performed, taxa were classified down to lowest available taxonomic classification. Species and environmental variables (depth [m], salinity, and average *Fucus* thalli width [cm], height [cm] and wet weight [g]) with a significance level set at 0.05 were included within the plot. *Fucus* morphological measurements were acquired from the open access dataset: 10.6084/M9.FIGSHARE.19690930 (Preston and Rodil 2022). We examined the habitat (fixed factor, *Fucus* vs Bare) dissimilarity of the macroinfauna assemblages (i.e. species-specific abundance) across countries (fixed, Sweden vs Finland) and sites (random factor, three levels nested in country) using non-parametric multivariate analyses of variance (PERMANOVA) based on the Bray-Curtis resemblance measure calculated from 4^th^-root transformed data (4999 unrestricted permutations, Type III). Epifauna assemblages were examined using only two factors (Country and sites). Changes in the macrofauna abundance (ind m^-2^) and the number of taxa were analysed through a 3 way (infauna) and a 2-way (epifauna) PERMANOVAs (same factors as above). We calculated distance resemblance matrices using Euclidean dissimilarity measures based on non-transformed data (4999 unrestricted permutations). For the eDNA analyses, a matrix with the relative abundance of each macroinfauna species (normalized by presence/absence) was used to examine the habitat dissimilarity of the assemblages across countries and sites (same as above) based on the Bray-Curtis resemblance measure. Changes in the number of species (log (x+1)-transformed eDNA asv reads) were analysed through a 3-way PERMANOVA (same as above) using Euclidean dissimilarity resemblance matrix. Only significant effects (p < 0.05) were further investigated by pairwise comparisons. PERMANOVA analyses were performed using PRIMER7 (Anderson et al. 2008). Abundance (individuals per m^2^ or sequences per site replicate) and species richness were plotted using ggplot2 (Wickham 2016) for the macroinfaunal (both habitats), epifaunal and eDNA samples. Additionally mean abundances (per m^2^) for the two conventional approach sampling techniques were combined per *Fucus* site and plotted alongside bare-sediment site abundances in ggplot2 (Wickham 2016). Shannon-Wiener diversity indices were calculated for the combined conventional approach and eDNA by using the vegan package (Oksanen et al. 2022) and plotted using ggplot2 (Wickham 2016).

## 3 Results

### 3:1 Comparison of methods

Totals of 41 and 130 separate taxa were detected through the combined conventional or the eDNA approaches respectively. Venn diagrams showed that there was variation between the ability of the approaches to capture biodiversity (Figure 2). However, the two approaches had considerable overlap in the classes detected (Askö 6, Tvärminne 4), although both approaches also detected exclusive taxa (eDNA: Askö 7, Tvärminne 8; conventional: Askö 1, Tvärminne 3). Overall the eDNA approach detected a larger amount of classes (Askö 17, Tvärminne 18) compared to the conventional approach (Askö 12, Tvärminne 10).

**Figure 2:**
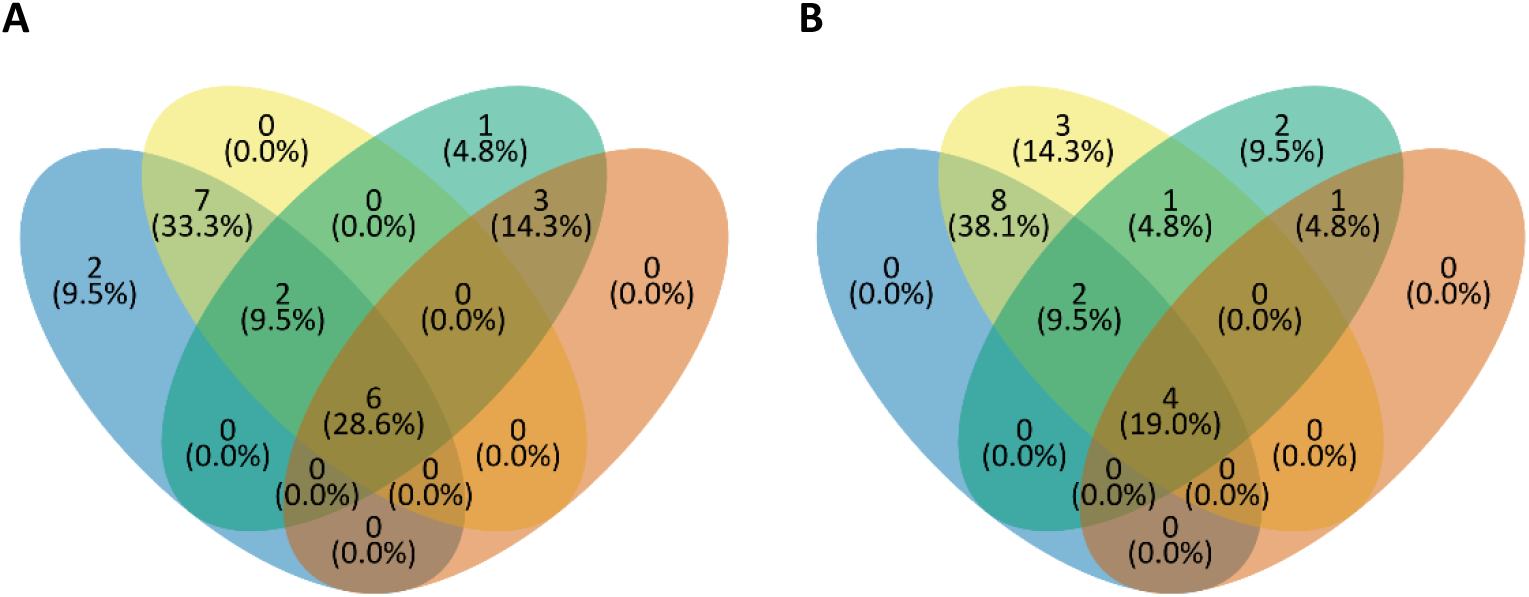
Proportion of classes detected by conventional vs eDNA approaches across habitats for Askö (A) and Tvärminne (B). Colour key: blue, bare-sediment eDNA; yellow, *Fucus* eDNA; green, *Fucus* conventional; orange, bare-sediment conventional.

### 3:2 Assemblage composition

The sampling approaches determined relatively dissimilar biodiversity estimates with a greater similarity seen among samples from each approach compared to between habitat types with different approaches (Figure 3A, B). The composition of assemblages detected by the conventional approach represented three major phyla (Annelida, Arthropoda, and Mollusca) whereas the eDNA approach represented four major phyla (Annelida, Arthropoda, Nematoda, and Platyhelminthes) with high levels of variability. Using the conventional approach, the most abundant taxa include: Annelida: *Marenzelleria* sp., *Hediste diversicolor*,and Oligochaeta; Arthropoda: various *Insecta* spp. (particularly *Chironomus* sp.), *Balanus* spp., and *Ostracoda* spp.; and Mollusca: *Macoma balthica, Mytilus* spp. complex, *Peringia ulvae*, and *Theodoxus fluviatilis* (Figure 3C, D, Supplementary material S2). The majority of these abundant taxa were not detected through the eDNA approach (Figure 3E). *Macoma balthica* and *P. ulvae* were detected by eDNA, although in relatively low numbers. The taxa with higher relative abundances detected by the eDNA approach included: Annelida: *Potamothrix* sp., *H. diversicolor;* Arthropoda: *Acartia tonsa, Candona bimucronata*, and *Semicytherura striata;* Mollusca: *Cerastoderma edule*, and Nematoda: *Sabatieria* sp. (Figure 3E, Supplementary material S3). Although the eDNA assemblages represented a high level of variability by site and habitat type. Overall the conventional approach presented a far simpler biodiversity estimate with less variability between sites/habitat. The approaches showed similar abilities to capture certain taxa (e.g. Arthropoda). Although relative abundances of Arthropoda in the eDNA dataset were in general far larger. Conversely, Mollusca were well represented by the conventional approach whilst being poorly represented or entirely absent from the majority of eDNA samples. Notability, a large proportion of the relative abundance captured by the eDNA approach was represented by taxa too small to be detected by the conventional approaches applied (e.g. Nematoda, Kinorhyncha, and Rotifera). No taxa were clearly confined to each habitat, irrelevant of the approach used (Supplementary material S2, S3).

**Figure 3:**
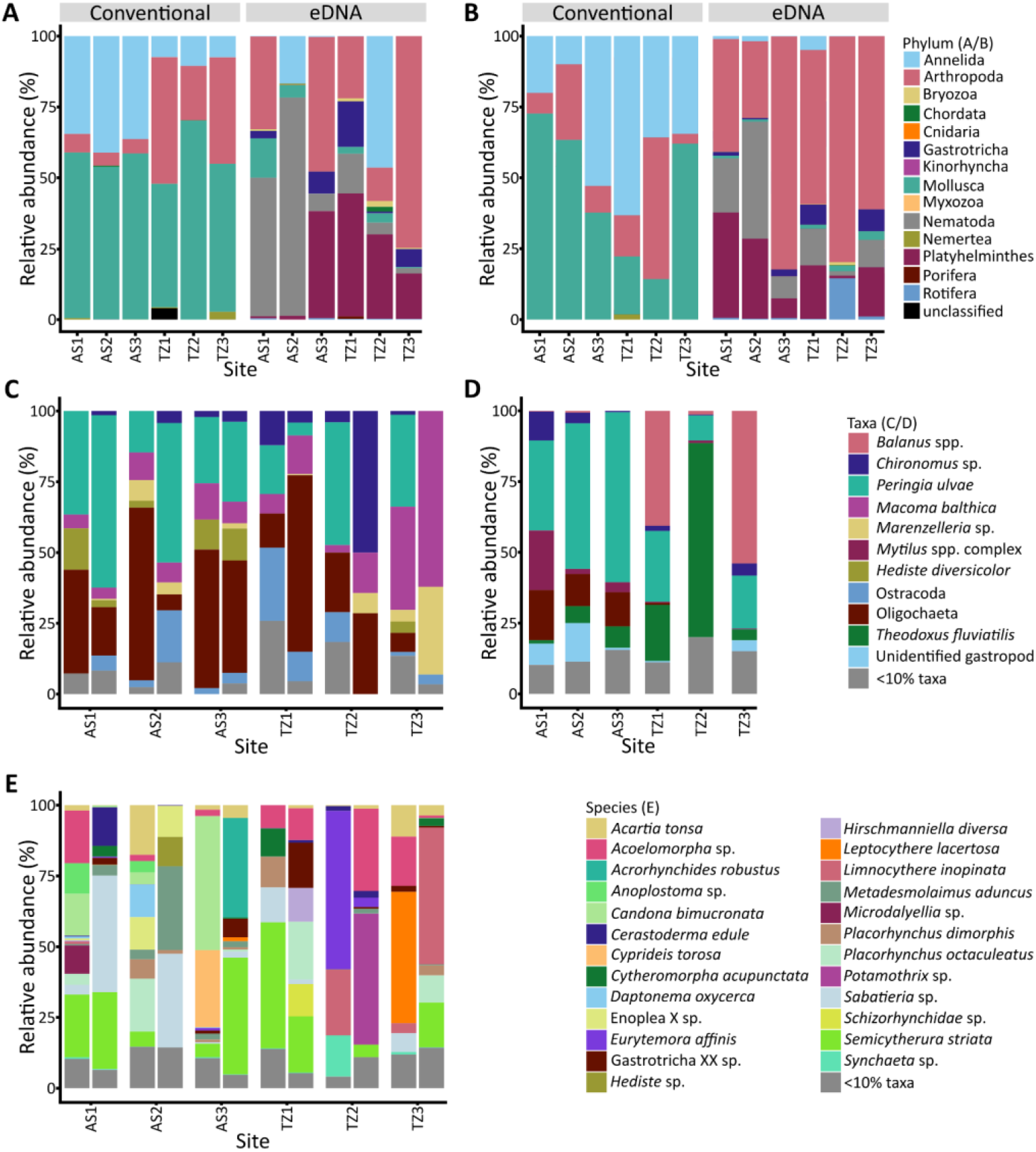
Phylum-level metazoan diversity of *Fucus* (A) and bare sediment (B) habitats when using the conventional or eDNA approaches. Diversities to the lowest available taxonomic ranking of macroinfaunal (C) and epifaunal (D) assemblages when using the conventional approach and eDNA approach (E). *Fucus* (left) and bare sediment (right) habitat diversities plotted side by side (C) and only for *Fucus* habitats (D). In plots C and D taxa with a relative abundance <10 % grouped into a single category.

The conventional approach allowed a more traditional description of community assemblages comparable to many previous studies (e.g. Rinne et al. 2022; Schagerström et al. 2014) (Figure 3C, D). Assemblages varied by site, country, and habitat although few discernible trends were observed. The macroinfauna and epifauna assemblages shared taxa (e.g. *Chironomus* sp., Oligochaeta, and *P. ulvae*) although exclusive taxa were also present. Both habitat type and country influenced the assemblages of the shared taxa, though trends were not uniform. Greater relative abundances of Oligochaeta within macroinfauna assemblages were associated with *Fucus* at Askö, whilst the reverse was true at Tvärminne. In opposition, greater abundances of *P. ulvae* within macroinfauna assemblages tended to be associated with bare sediments at Askö and *Fucus* habitats in Tvärminne. However *P. ulvae* were fairly abundant in all epifaunal samples, particularly at Askö. *Chironomus* sp. were more associated with bare habitats in the macroinfauna assemblages at Askö, however *Chironomus* sp. were relatively common in the epifaunal samples from two sites at Askö (AS1, AS2). *Macoma balthica* were present within the macroinfauna of all sites, being particularly well represented in TZ3. The abundance in relation to habitat type was similar to that observed in Oligochaeta, whereby *M. balthica* were more associated with *Fucus* at Askö and bare sediments at Tvärminne. Overall, the conventional datasets demonstrated a high level of variability among assemblages at the habitat, country and site level.

MRPP ordination showed that sites associated with *Fucus* generally were ordinated closer to those from the same country in both traditional sampling communities, though all sites were fairly similar (Figure 4). The eDNA dataset showed particularly close ordination with high similarity between all sites. PERMANOVAs supported the trend of similarity between countries, showing no significant differences by country (macroinfauna: Pseudo-F = 2.38, P = 0.103; epifauna: Pseudo-F = 3.65, P = 0.103; eDNA: Pseudo-F = 1.36, P = 0.304; Supplementary material S4, S5). Although within country assemblages varied significantly by site (macroinfauna: Pseudo-F = 3.00, P = 0.0001; epifauna: Pseudo-F = 3.69, P = 0.0001; eDNA: Pseudo-F =3.80, P = 0.0001; Supplementary material S4, S5). Within each country the number of exclusive taxa ranged from 17.1-36.9% of the observed taxa depending on the approach (Supplementary material S6, S7). In the traditional dataset, the ordination of the two countries is partially influenced by a group of significant species, particularly in the case of epifauna assemblages (Figure 4). Sites at Tvärminne were more associated with *Balanus* spp., *Potamopyrgus antipodarum, T. fluviatilis*, and Zygoptera, whilst sites at Askö associated with *Diptera* sp., *Chironomus* sp., *Peringia ulvae, Mytilus* spp. complex, Oligochaeta, and *Parvicardium spp.*. Taxa significantly influencing the ordination of eDNA were Enoplea, *Hediste* sp., *Metadesmolaimus aduncus*, and *Parvicardium spp*. The environmental variables of depth and one morphological variable (either thallus height or wet weight) significantly influenced the ordination of the traditional samples (Figure 4). Environmental variables did not influence the ordination of the eDNA plot.

**Figure 4:**
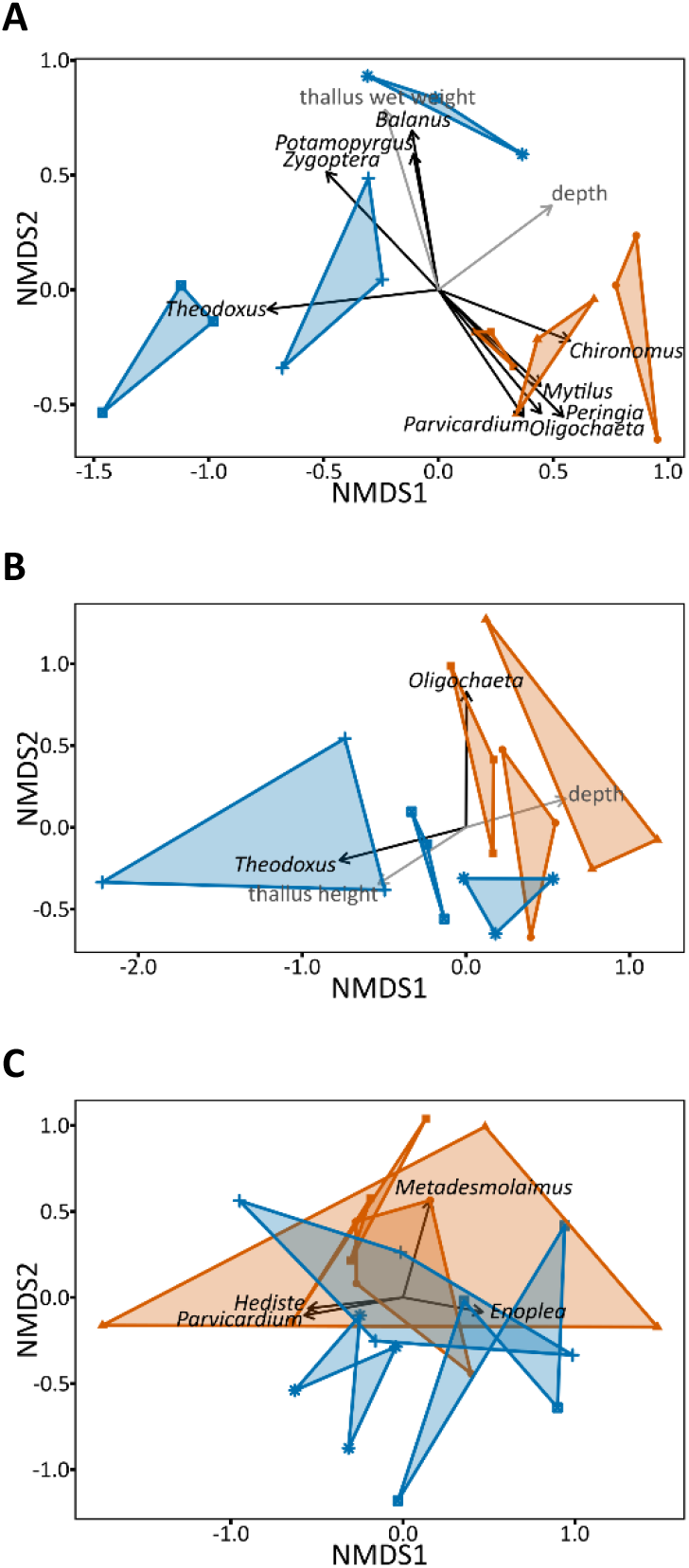
Multiple Response Permutation Procedure (MRPP) ordination of taxa-specific abundance of the epifaunal (A), macroinfaunal (B), and eDNA (C) benthic communities of free-living *Fucus*. Three (A, B) or four (C) samples per site are shown. Coordinates are shown as Non-Metric Multidimesional Scaling (NMDS) ordination, based on the dissimilarity matrix between sites. Site abbreviation/colours: AS/orange, Askö; TZ/blue, Tvärminne. Site symbols: ●, AS1; ▲, AS2; ■, AS3; +, TZ1; ⊠, TZ2; ✳, TZ3.

### 3:3 Biodiversity estimates

Using the conventional approach, *Fucus* habitats determined lower species richness and abundance of macroinfauna assemblages compared to the adjacent bare-sediment habitats in four sites (Askö [3], Tvärminne [1]) (Figure 5). Conversely, the remaining two sites showed far greater species richness and abundance in the *Fucus* associated habitat (Tvärminne [2]). Consequently, *Fucus* cannot be considered to have a uniform influence on macroinfauna assemblages. PERMANOVAs determined that habitat type had varying influence on the composition of assemblages. Within the eDNA dataset, the composition of assemblages was significantly affected by the type of habitat within Askö, yet habitat had no significant effect within Tvärminne (Askö: t = 1.85; P = 0.017; Tvärminne: t = 1.08; P = 0.399; Supplementary material S5). Similarly, a few sites in the conventional macroinfaunal dataset displayed significantly different assemblages based on habitat type (Supplementary material S5). *Fucus* habitats were associated with additional epifauna assemblages, however the epifaunal abundances were much lower than the abundances of the macroinfauna assemblages associated with bare sediments at four of the sites (AS1-3, TZ1) (Figure 5). Species richness, on the other hand, was greater in epifauna assemblages. When considering combined macroinfauna and epifauna assemblages (Figure 6A, B), four sites had greater abundances associated with the *Fucus* habitat, but abundances in the two other sites were nearly doubled in the bare-sediment compared to the neighbouring *Fucus* habitat. Thus, no uniform trend in abundances can be seen in the combined faunal communities relating to the presence of *Fucus*, with influences being site specific. However *Fucus* has a positive influence on the number of species recorded, with the presence of *Fucus* correlating to an at least doubled species richness compared to the bare sediment. Similar to the conventional approach, eDNA does not show a uniform pattern in the influence of *Fucus* on biodiversity (Figure 6C, D). The mean sequence abundance was higher within the *Fucus* habitat for four sites (Askö [2], Tvärminne [2]) whilst mean species richness was higher in three (Tvärminne [3]). Although in all but one of these sites, the means are relatively similar to the neighbouring bare sediment means. Of the two sites with lower abundances associated with *Fucus* habitats, the means are much below the bare-sediment means. Species richness was higher in *Fucus* habitats compared to bare sediment habitats at Tvärminne whilst the opposite was true at Askö. Species diversities, in the form of the Shannon index, contrast by sampling approach (Figure 7). Using the conventional approach species diversity is greater in soft sediments associated with *Fucus* whilst the eDNA approach illustrates no link between species diversity and presence/absence of *Fucus*.

**Figure 5:**
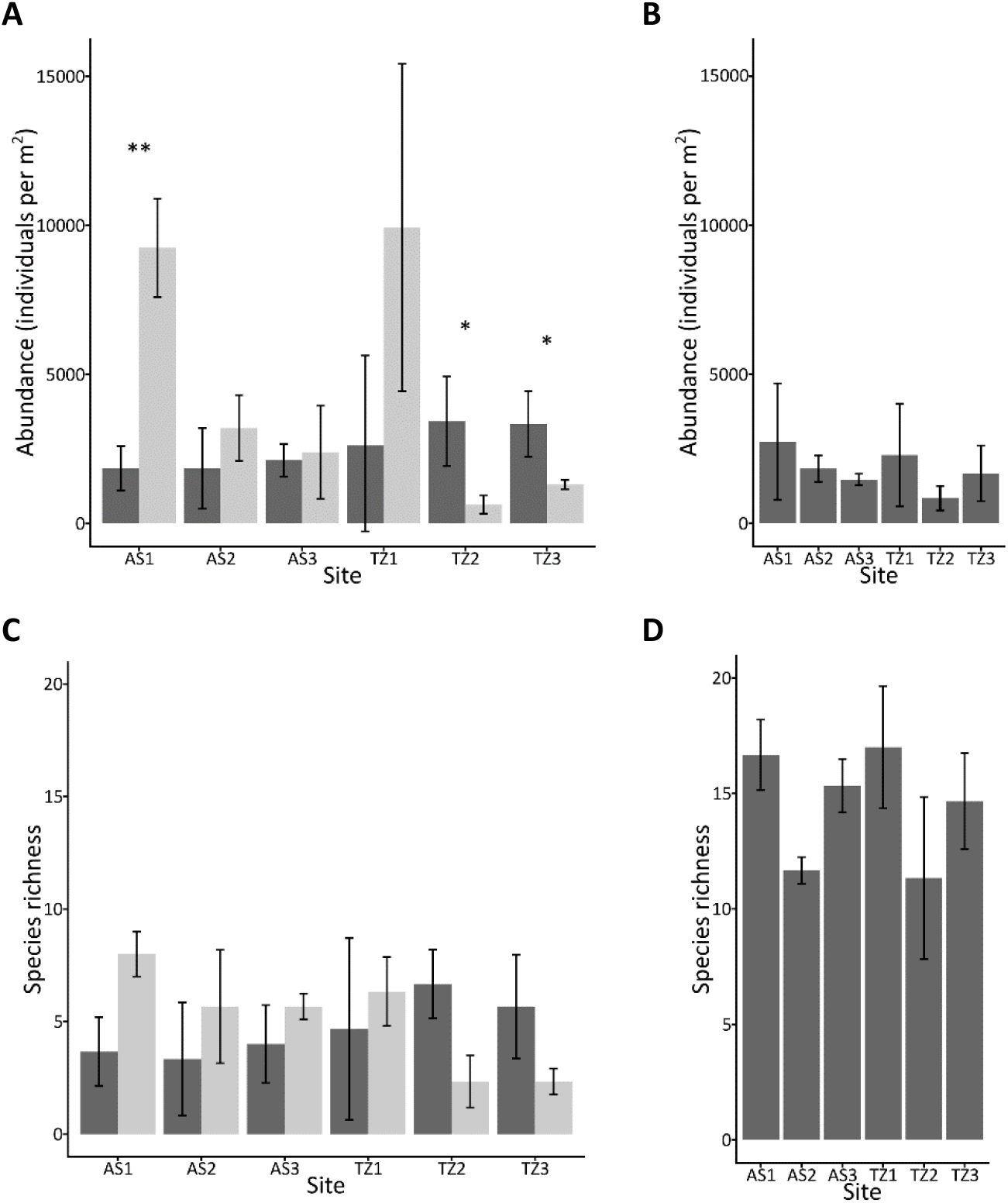
Total abundance (A, B) and species (taxa) richness (C, D) of macroinfauna (A, C) and epifauna (B, D) in samples from sites with and without free-living *Fucus* in Askö and Tvärminne using the conventional approaches. Abbreviations: AS, Askö; TZ, Tvärminne. Colour codes: Dark grey, *Fucus* habitats; light grey, bare-sediment habitats. Significance levels of PERMANOVA post hoc tests between habitat types within sites.

**Figure 6:**
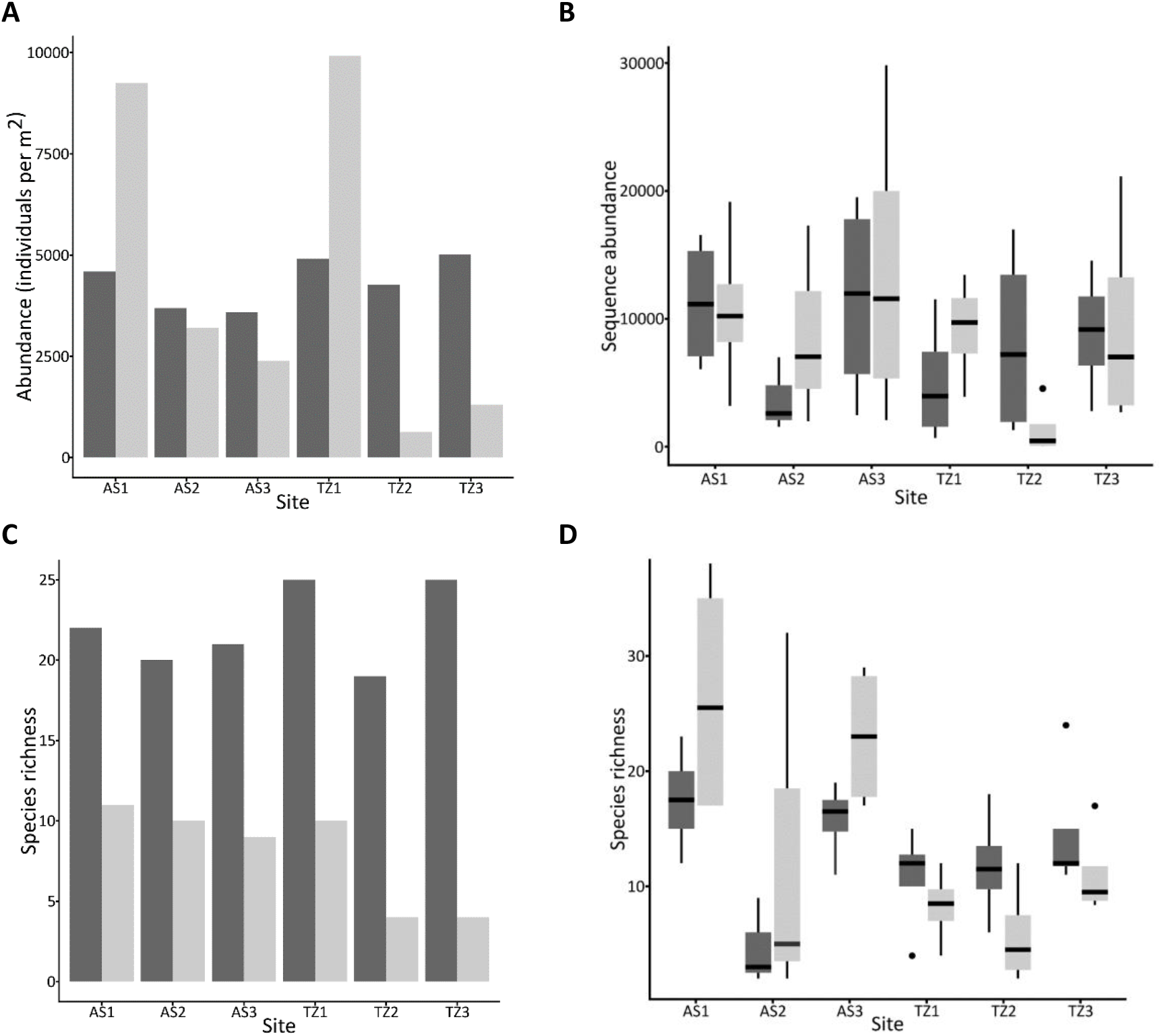
Mean total abundance (A) and total species richness (C) of the macroinfauna and epifauna communities combined from sites with and without free-living *Fucus* in Askö and Tvärminne. Sequence abundance (B) and species richness (D) of benthic communities from sites with and without free-living *Fucus* in Askö and Tvärminne using eDNA. Abbreviations: AS, Askö; TZ, Tvärminne. Colour codes: Dark grey, *Fucus* habitats; light grey, bare-sediment habitats.

**Figure 7:**
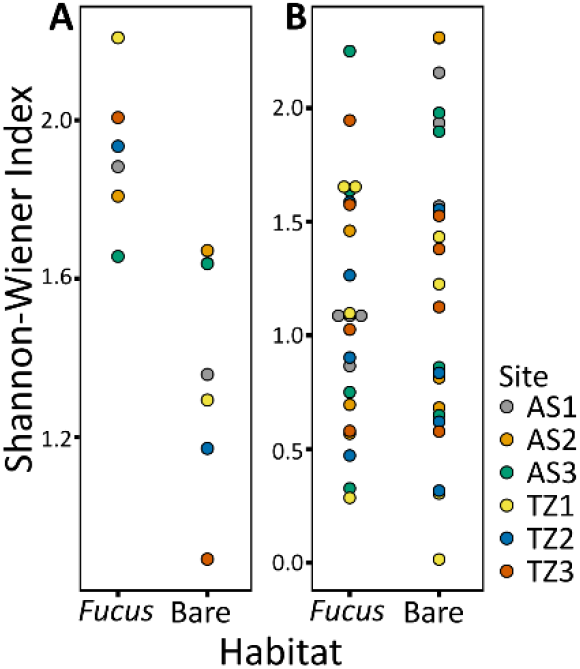
Shannon-Wiener diversity index of habitat types by sampling approach: conventional (A), eDNA (B). Abbreviations: AS, Askö; TZ, Tvärminne.

## 4 Discussion

In this study we show that (i) *Fucus* had varying influence on the faunal benthic assemblages but appeared to increase species richness in many of the sites, and (ii) biodiversity estimates varied depending on the approach employed.

Biodiversity estimates were highly variable between the sampling sites, with few consistent trends being observed within and between sites and habitat types. We surmise that this is due to varying abiotic and biotic factors among the sites. For example, benthic communities are known to differ by depth (Orav et al. 2000), oxygen concentration (Lauringson and Kotta 2006), eutrophication (Rinne et al. 2022), sediment type (Kotta and Orav 2001; Mosbahi et al. 2016), thickness of algal cover (Lauringson and Kotta 2006), algae thalli size (Schagerström et al. 2014), and algae thalli structural complexity (Cacabelos et al. 2010; Hansen et al. 2011; Kraufvelin et al. 2006). The free-living *Fucus* sites within our study differ abiotically, including in the recorded variables of salinity (5.9–6.1) and depth (max 1.9–3.4 m), but also in the morphotypes present at each site (Preston and Rodil 2023) which may explain the lack of congruence. In the conventional approach MRPP, depth was seen to have a significant influence and the sites with similar max depths grouped more closely (e.g. AS2/AS3 and TZ1/TZ2). Due to the often small patch size (~10–20 m^2^) of free-living populations on mostly gently sloping, bottom gradients and the current poor understanding of their distribution, it is challenging to select sites to mitigate the influence of these varying abiotic conditions. Consequently we highlight that our free-living sites were heterogeneous in nature.

Nevertheless when considering all sites using the conventional approach and Tvärminne using the eDNA approach, species richness was positively associated with *Fucus* as a result of additional epifauna assemblages. Higher Shannon diversity was also observed in the *Fucus* sites of the conventional approach. Therefore we suggest that the presence of *Fucus* can positively influence biodiversity through creating a complex habitat which increases the number of niches and consequently supporting more species (Kostylev et al. 2005; Levin 1992; Pianka 2011). Similar trends have been observed within the Baltic Sea for various macrophyte communities including *Zostera marina* (Boström and Bonsdorff 1997), *Furcellaria lumbricalis* (Kotta and Orav 2001), and drift algae mats (Lauringson and Kotta 2006). However, using eDNA this trend is not as apparent, with the mean species richness being higher within the three *Fucus* habitats at Tvärminne but lower within all sites at Askö compared to the bare-sediment habitats. Shannon diversities were also highly variable with no apparent trend relating to habitat type.

*Fucus* constrains the species richness and abundance of macroinfauna assemblages in four of the six sites. In fact, the bare-sediment habitats of these four sites showed far greater macroinfaunal abundances (e.g. a 5-fold difference at AS1) compared to the neighbouring *Fucus* habitat. A comparable macroalgae, *F. lumbricalis*, which also forms free-living populations throughout the Baltic Sea, has been found to demonstrate a similar trend of reducing macrozoobenthos (Kotta and Orav 2001). However within attached *Fucus* canopies, abundance has been found to be higher in poor quality habitats despite the lower observed species diversity (Rinne et al. 2022). Consequently the relationship between algae habitat-formation and patterns of biodiversity is complex and partially depends on the biodiversity metric used. It appears that the presence of *Fucus* often has a filtering effect, exerting exclusionary pressure on organisms with traits suited to bare sediments. Overall, *Fucus* appears to frequently be detrimental to macroinfauna assemblages yet is able to alleviate some of the negative effects through the provision of habitat usable by epifaunal organisms. Although this trend is not universal since in two sites (TZ2, TZ3) *Fucus* greatly improves species richness and abundance of both the macroinfauna and epifauna assemblages.

Three phyla (Annelida, Arthropoda, and Mollusca) were well represented in the conventional dataset, whilst four phyla (Annelida, Arthropoda, Nematoda, and Platyhelminthes) were well represented in the eDNA dataset. Nematoda and Platyhelminthes were not detected using the conventional approach due to their small size as meiobenthos, predisposing these phyla against detection using the selected conventional approaches (Ojaveer et al. 2010). Mollusca were poorly represented by the eDNA approach despite being abundant within the conventional dataset. In the eDNA approach, the non-detection of taxa does not automatically imply its absence, equally a positive-detection does not necessarily indicate that the taxa is currently present because the eDNA could have been transported over space or preserved over time (Roussel et al. 2015). In fact, false positives and negatives are common in eDNA (Beng and Corlett 2020). Further limitations to eDNA detection include the highly variable degradation rates of eDNA (Barnes et al. 2014; Beng and Corlett 2020), primer bias (Stadhouders et al. 2010), the potential for PCR inhibition (Harper et al. 2019; Jane et al. 2015; Schrader et al. 2012), and the quality of the reference database (Foster et al. 2022). Consequently, we infer that the failure to detect higher abundances of Mollusca are an artefact of the sampling approach rather than their absence within the environment. The aforementioned limitations may also contribute to the greater observed relative abundance of Arthropoda in eDNA, as the PCR conditions and reference database may favour their detection. To illustrate these potential discrepancies in greater detail, both *Chironomus* sp. and *Marenzelleria* sp. were abundant within the conventional dataset yet absent from the eDNA dataset. However, mismatches on the reverse primer for *Marenzelleria arctia* sequence suggest that *Marenzelleria* spp. were present yet limitations in the eDNA protocol hindered their detection. Likewise, the Baltic Sea *Chironomus* species (*Chironomus plumosus*) is known to represent a 450 bp long fragment while the sequence reads within this study were max 401 bp. Consequently *Chironomus* spp. reads were likely to be in the dataset but forward and reverse reads did not merge, hence the apparent absence. The discordant limitations of each approach resulted in the two sampling approaches illustrating differing biodiversity estimates, showing far greater similarity among biodiversity estimates within each approach irrelevant of habitat type, site and country. Thus the use of either approach favours the selection of certain taxa whilst hindering others. Consequently the two approaches are not directly comparable, yet the combined use can provide new insights into the biodiversity supported by these habitats at diverse scales. Although biodiversity estimates diverge by approach, both support similar assemblages between the two habitats suggesting that the presence of *Fucus* does not exert extreme habitat filtering effects. Overall, our study highlights the necessity to consider the limitations of sampling approaches when generating biodiversity estimates; but also that biodiversity estimates can benefit greatly from a multi-approach design where both conventional and eDNA approaches are employed in complement.

This study provides new insights into the associated assemblages of free-living *Fucus*. Free-living *Fucus* supports similarly diverse macrofauna assemblages to the attached form, with abundant taxa including *Chironomus* sp., Oligochaeta, *P. ulvae*, and *T. fluviatilis* (Rinne et al. 2022; Schagerström et al. 2014). *Idotea* spp., a genus commonly associated with attached *Fucus* (Korpinen et al. 2007; Schagerström et al. 2014) were poorly detected through either approach, although *Idotea* spp. have also in cases been observed in very low abundances on attached *Fucus* (Rinne et al. 2022). Thus the absence of *Idotea* spp. may relate to stochastic changes (Engkvist et al. 2000; Kangas et al. 1982) or directly to the characteristics of the free-living form. Patterns of assemblages were highly variable, at both the small (site) and large (country) spatial scale, although this variability was not significant between countries. Several exclusive taxa were observed for each country, as was expected due to the previously determined spatial patterns of benthic diversity (Zettler et al. 2014). We suggest that, similarly to the attached form, different abiotic and biotic drivers shape the associated assemblages (Rinne et al. 2022).

## Author contributions

RP: Conceptualization (lead), Formal analysis (lead), Funding acquisition (lead), Investigation (equal), Methodology (lead), Project administration (lead), Visualization (lead), Writing - Original Draft (lead). MM: Formal analysis (equal), Writing - Review & Editing (equal). VH: Formal analysis (supporting), Funding acquisition (supporting), Investigation (equal), Writing - Review & Editing (supporting). IFR: Conceptualization (supporting), Investigation (supporting), Methodology (supporting), Writing - Review & Editing (equal).

## Acknowledgments

We are grateful to Susanne Qvarfordt, Oskar Nyberg, and Chiara D’Agata for their assistance with fieldwork at Askö Laboratory.

## Funding information

Funding for this project was provided through grants from the Walter and Andrée de Nottbeck Foundation, the Onni Talas Foundation, and the Baltic Sea Centre (Askö grants).

## Conflict of interest

The authors declare that they have no known competing financial interests or personal relationships that could have appeared to influence the work reported in this paper.

## Data availability statement

Conventional and metabarcoding biodiversity data are available on Figshare: 10.6084/M9.FIGSHARE.20472768

## SUPPLEMENTARY MATERIAL

**S1:**
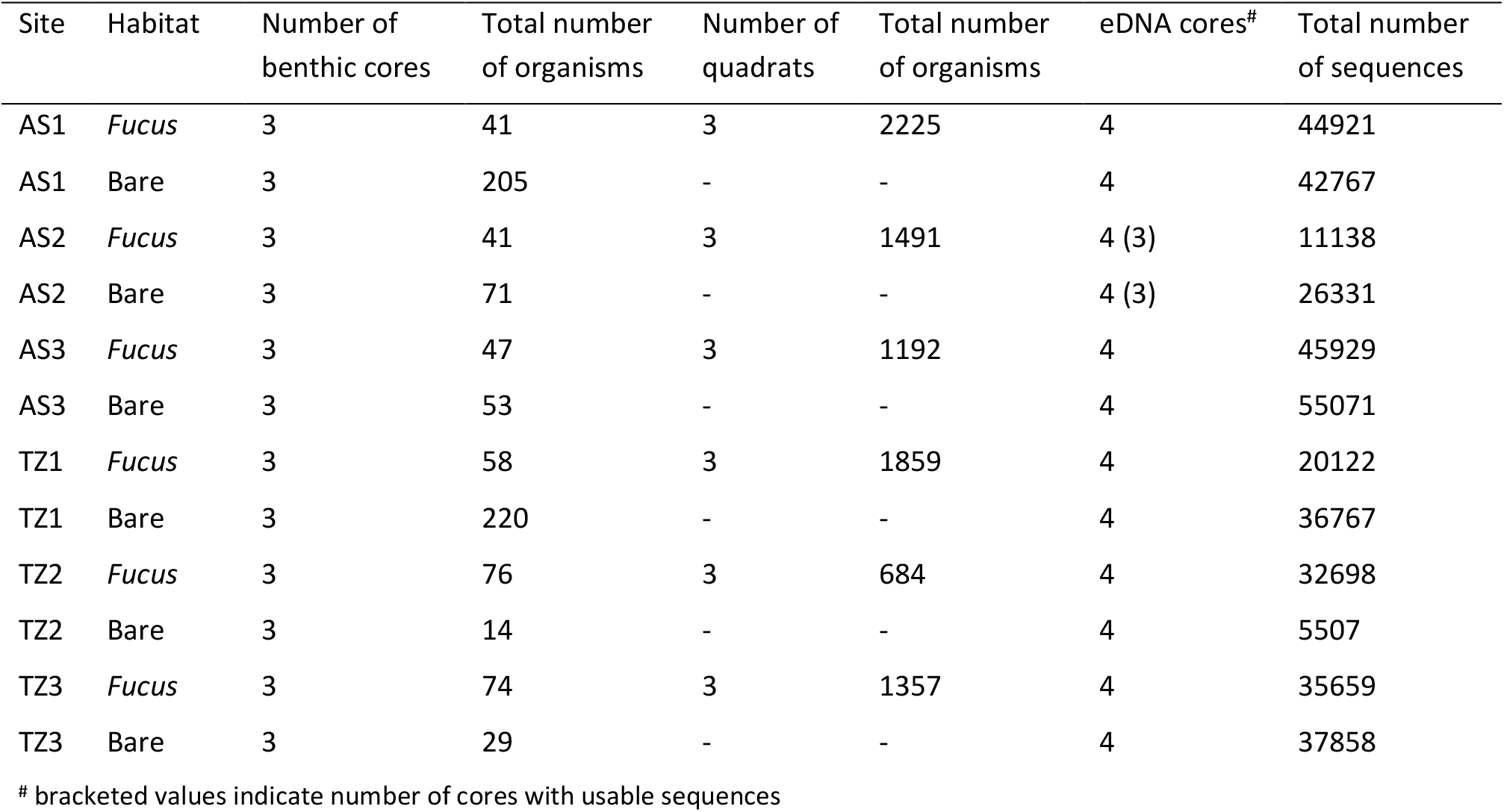
Field sampling information

**S2:**
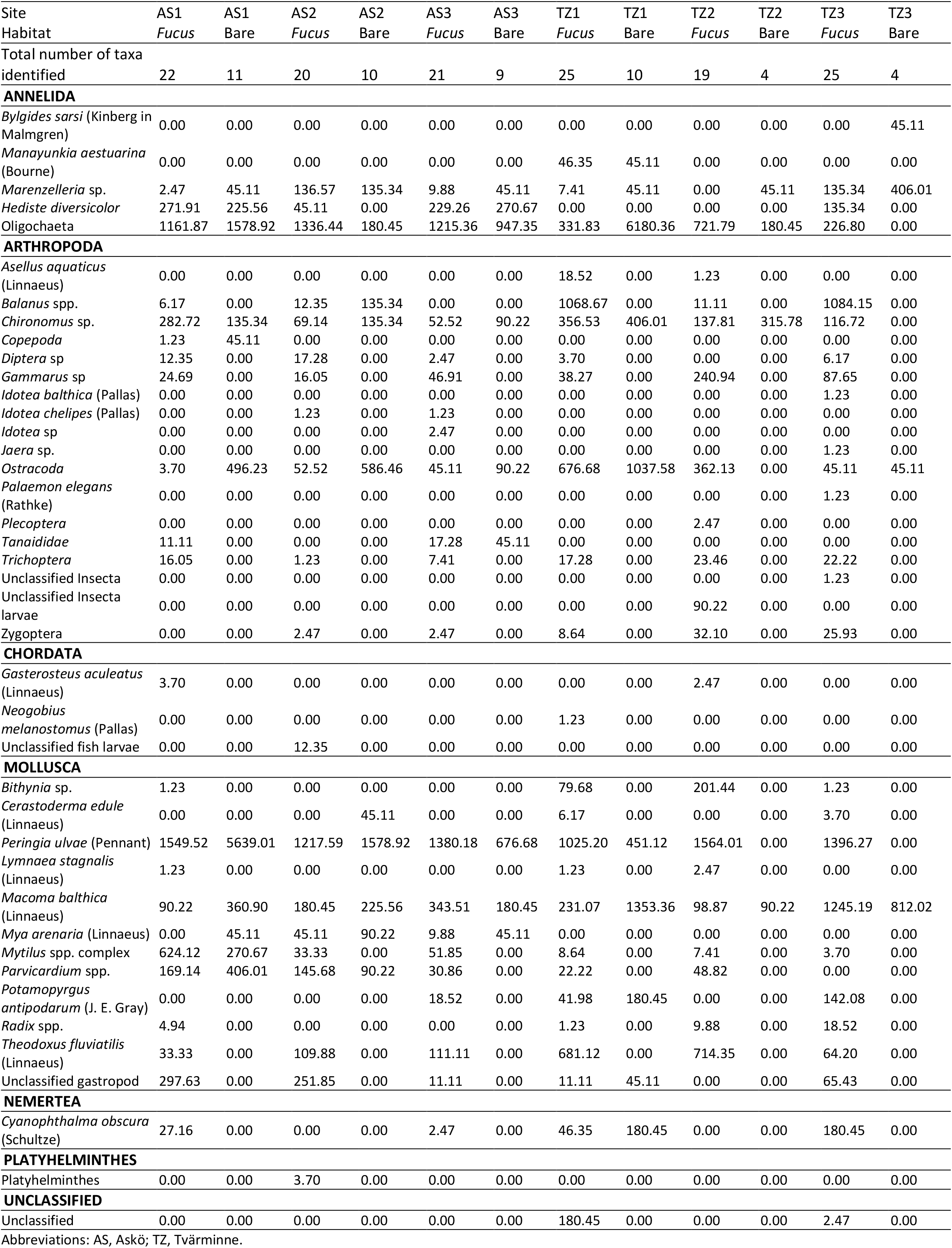
Combined macroinfauna and epifauna abundances per m^2^ from sites with and without free-living *Fucus* from Askö (Sweden) and Tvärminne (Finland) using the conventional approach.

**S3:**
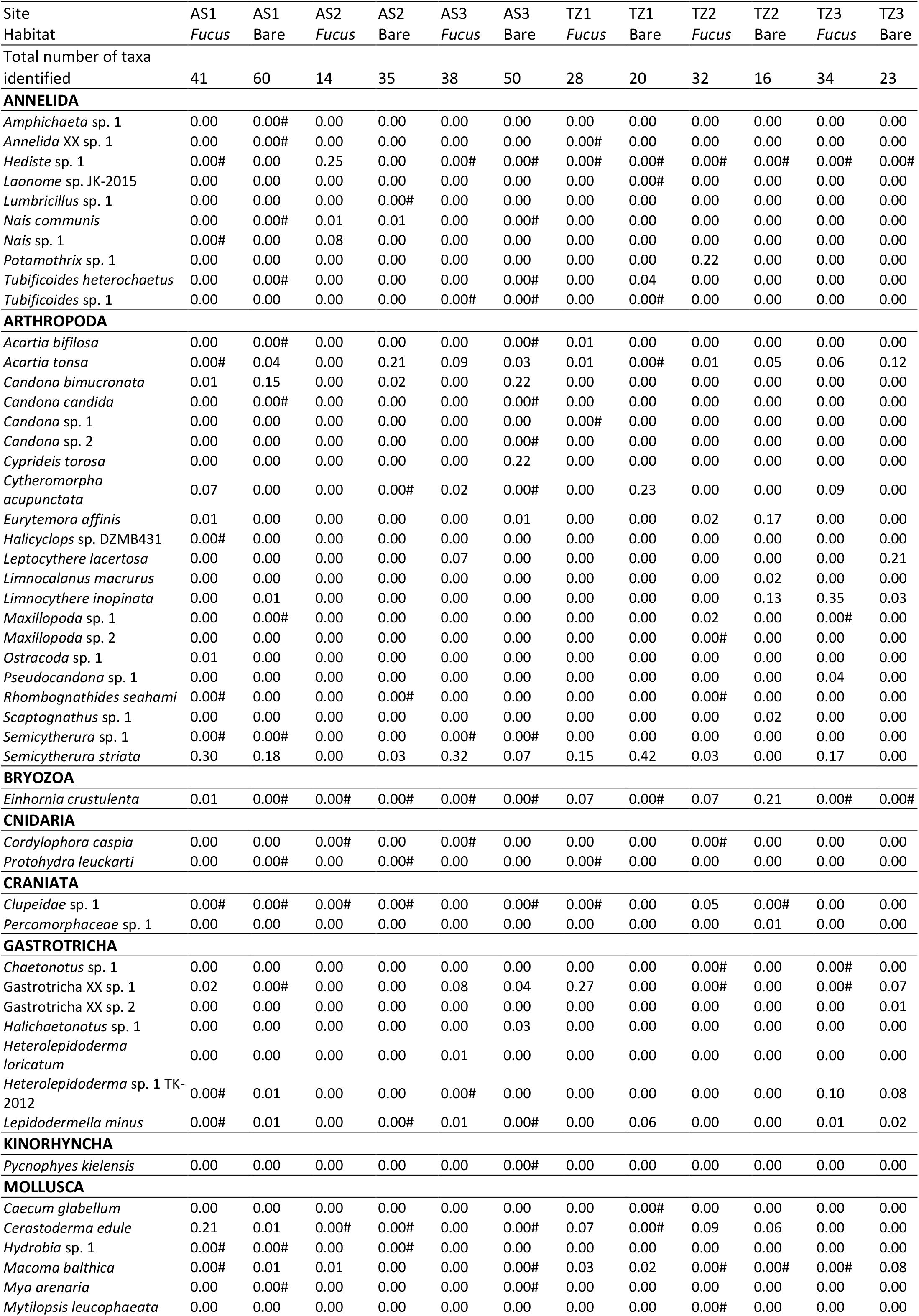
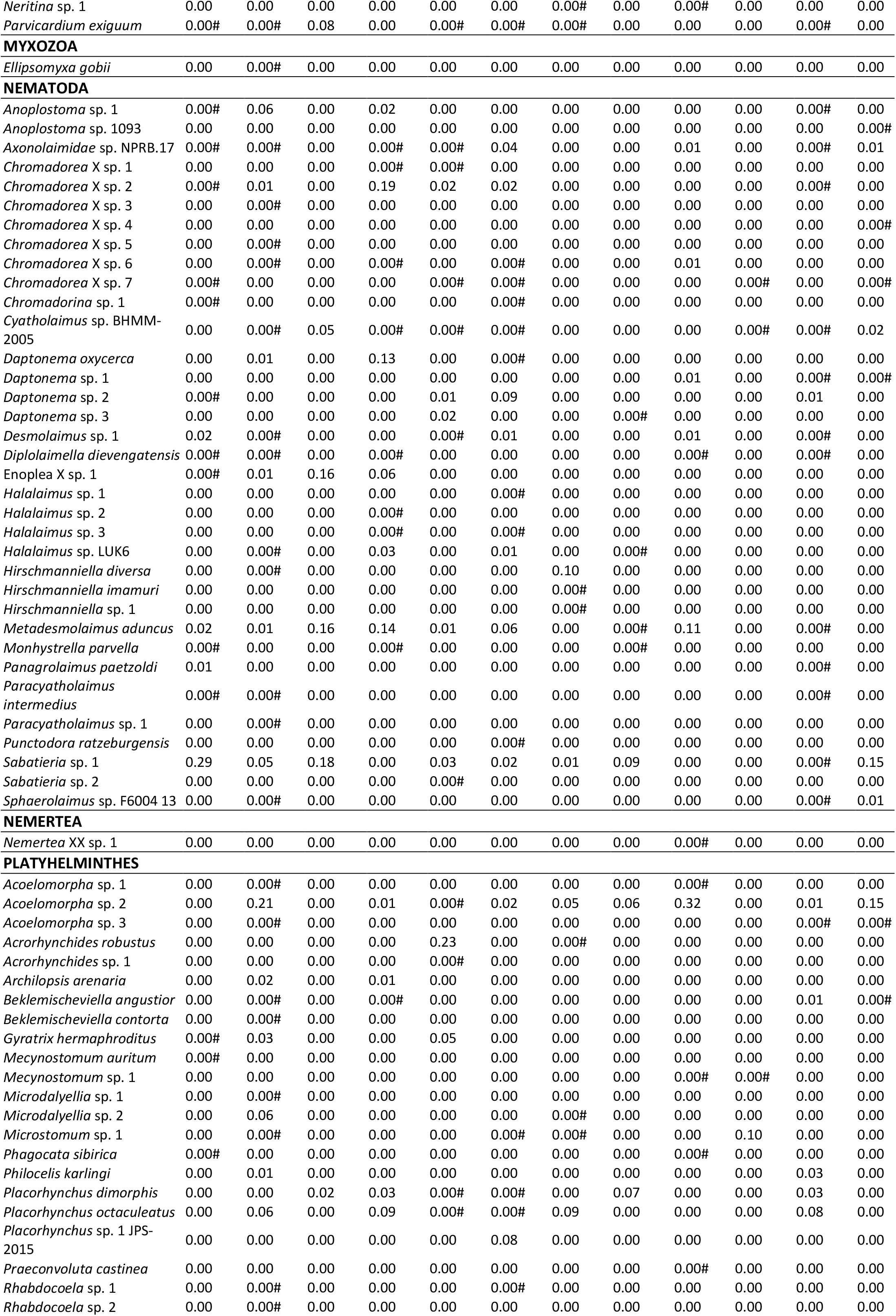
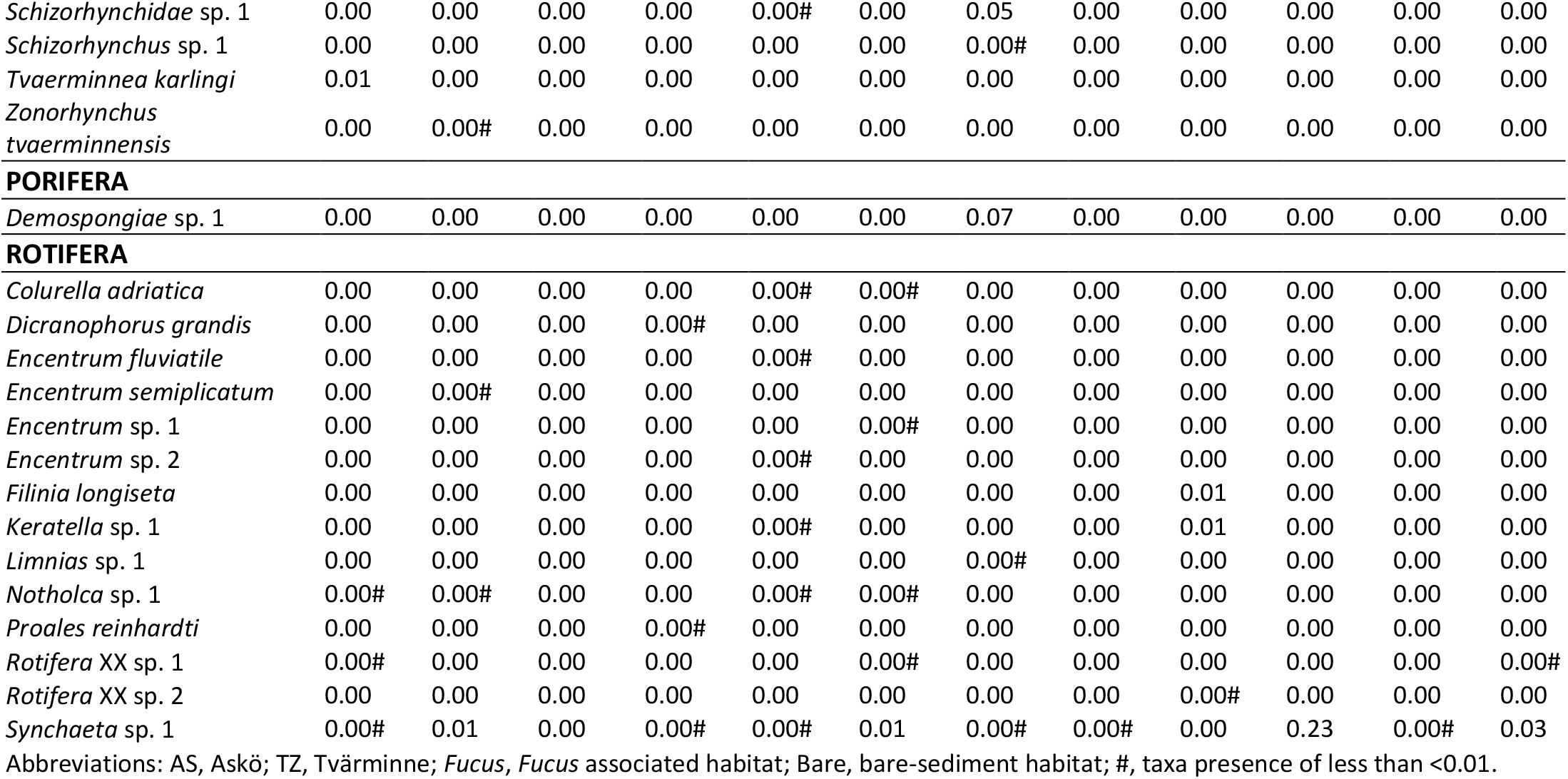
Relative abundances of metazoan assemblages from sites with and without free-living *Fucus* at Askö (Sweden) and Tvärminne (Finland) using the eDNA approach. Taxonomic affiliations from NCBI GenBank (23/05/22).

**S4:**
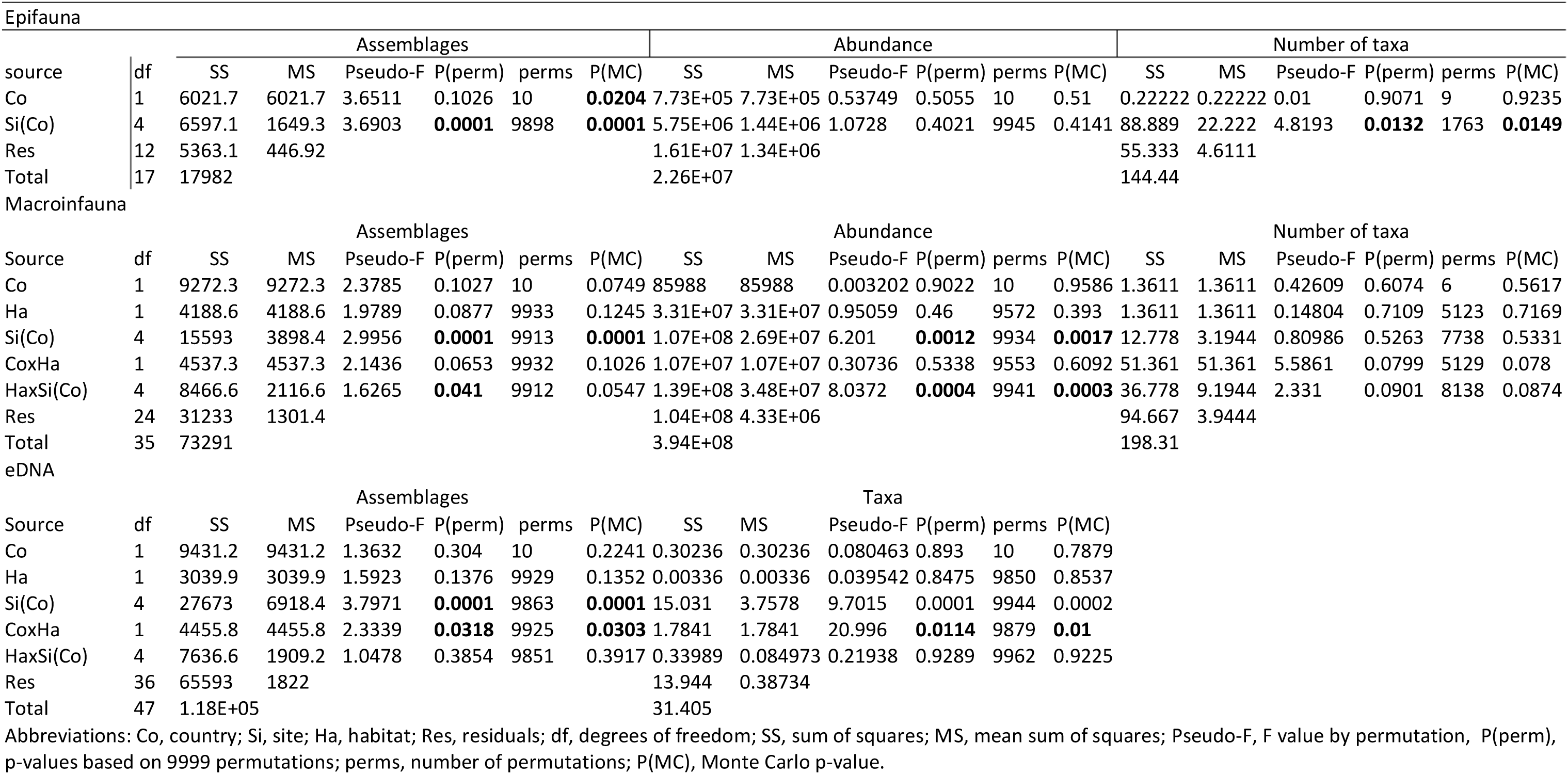
Permutational analysis of variance (PERMANOVA) results table for epifauna, macroinfauna, and eDNA. Country and Habitat as fixed factors and Site as random factor. Significance (p < 0.05) indicated in bold.

**S5:**
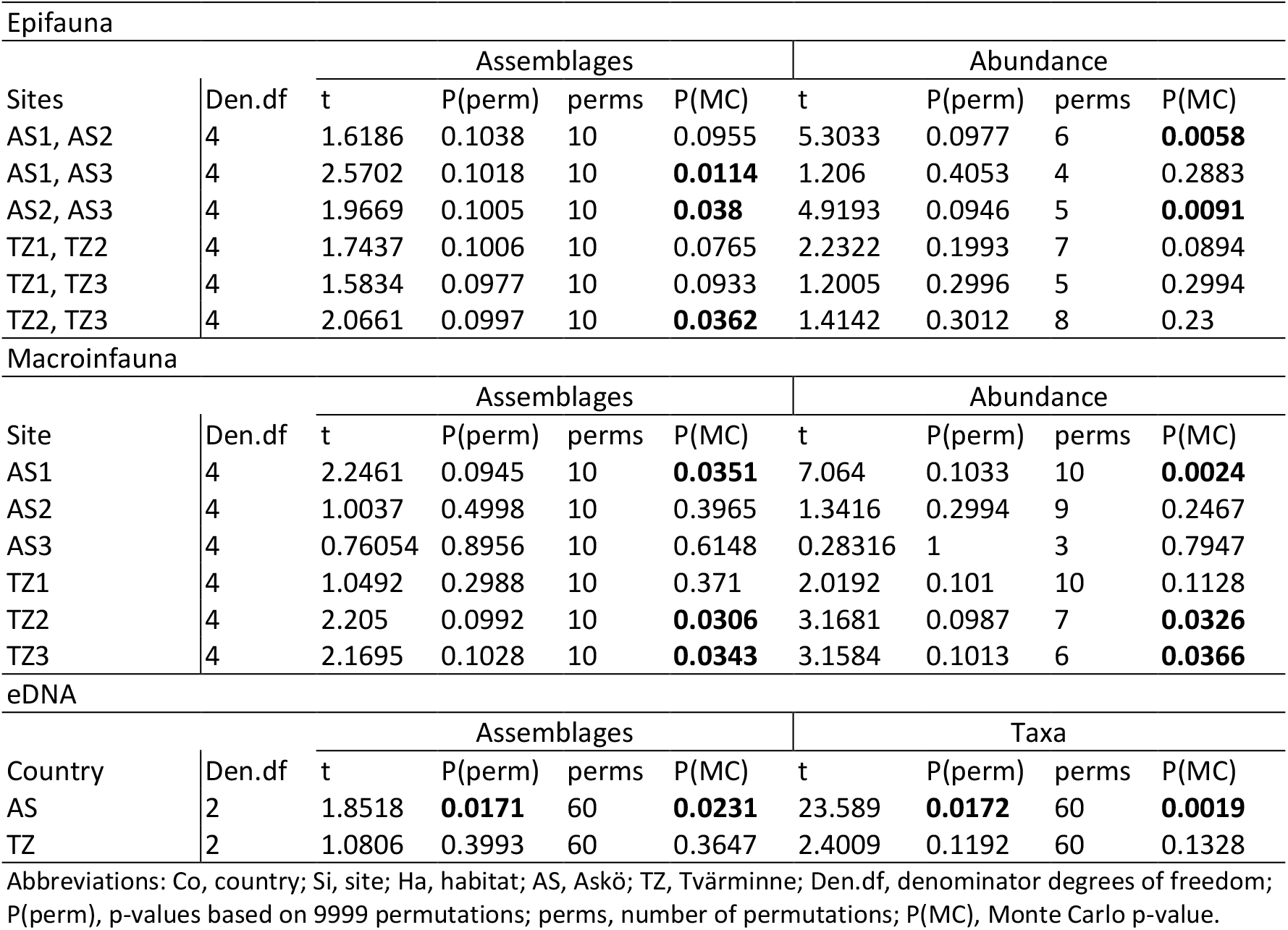
Post hoc tests for permutational analysis of variance (PERMANOVA) [S1]. Pair-wise tests: Epifauna, SI(Co); Macroinfauna, HaxSi(Co); eDNA, CoxHa.

| Epifauna |  |  |  |  |  |  |  |  |  |
| --- | --- | --- | --- | --- | --- | --- | --- | --- | --- |
| Sites | Den.df | Assemblages |  |  |  | Abundance |  |  |  |
|  |  | t | P(perm) | perms | P(MC) | t | P(perm) | perms | P(MC) |
| AS1, AS2 | 4 | 1.6186 | 0.1038 | 10 | 0.0955 | 5.3033 | 0.0977 | 6 | **0.0058** |
| AS1, AS3 | 4 | 2.5702 | 0.1018 | 10 | **0.0114** | 1.206 | 0.4053 | 4 | 0.2883 |
| AS2, AS3 | 4 | 1.9669 | 0.1005 | 10 | **0.038** | 4.9193 | 0.0946 | 5 | **0.0091** |
| TZ1, TZ2 | 4 | 1.7437 | 0.1006 | 10 | 0.0765 | 2.2322 | 0.1993 | 7 | 0.0894 |
| TZ1, TZ3 | 4 | 1.5834 | 0.0977 | 10 | 0.0933 | 1.2005 | 0.2996 | 5 | 0.2994 |
| TZ2, TZ3 | 4 | 2.0661 | 0.0997 | 10 | **0.0362** | 1.4142 | 0.3012 | 8 | 0.23 |

| Macroinfauna |  |  |  |  |  |  |  |  |  |
| --- | --- | --- | --- | --- | --- | --- | --- | --- | --- |
| Site | Den.df | Assemblages |  |  |  | Abundance |  |  |  |
|  |  | t | P(perm) | perms | P(MC) | t | P(perm) | perms | P(MC) |
| AS1 | 4 | 2.2461 | 0.0945 | 10 | **0.0351** | 7.064 | 0.1033 | 10 | **0.0024** |
| AS2 | 4 | 1.0037 | 0.4998 | 10 | 0.3965 | 1.3416 | 0.2994 | 9 | 0.2467 |
| AS3 | 4 | 0.76054 | 0.8956 | 10 | 0.6148 | 0.28316 | 1 | 3 | 0.7947 |
| TZ1 | 4 | 1.0492 | 0.2988 | 10 | 0.371 | 2.0192 | 0.101 | 10 | 0.1128 |
| TZ2 | 4 | 2.205 | 0.0992 | 10 | **0.0306** | 3.1681 | 0.0987 | 7 | **0.0326** |
| TZ3 | 4 | 2.1695 | 0.1028 | 10 | **0.0343** | 3.1584 | 0.1013 | 6 | **0.0366** |

| eDNA |  |  |  |  |  |  |  |  |  |
| --- | --- | --- | --- | --- | --- | --- | --- | --- | --- |
| Country | Den.df | Assemblages |  |  |  | Taxa |  |  |  |
|  |  | t | P(perm) | perms | P(MC) | t | P(perm) | perms | P(MC) |
| AS | 2 | 1.8518 | **0.0171** | 60 | **0.0231** | 23.589 | **0.0172** | 60 | **0.0019** |
| TZ | 2 | 1.0806 | 0.3993 | 60 | 0.3647 | 2.4009 | 0.1192 | 60 | 0.1328 |

Abbreviations: Co, country; Si, site; Ha, habitat; AS, Askö; TZ, Tvärminne; Den.df, denominator degrees of freedom; P(perm), p-values based on 9999 permutations; perms, number of permutations; P(MC), Monte Carlo p-value.

**S6:**
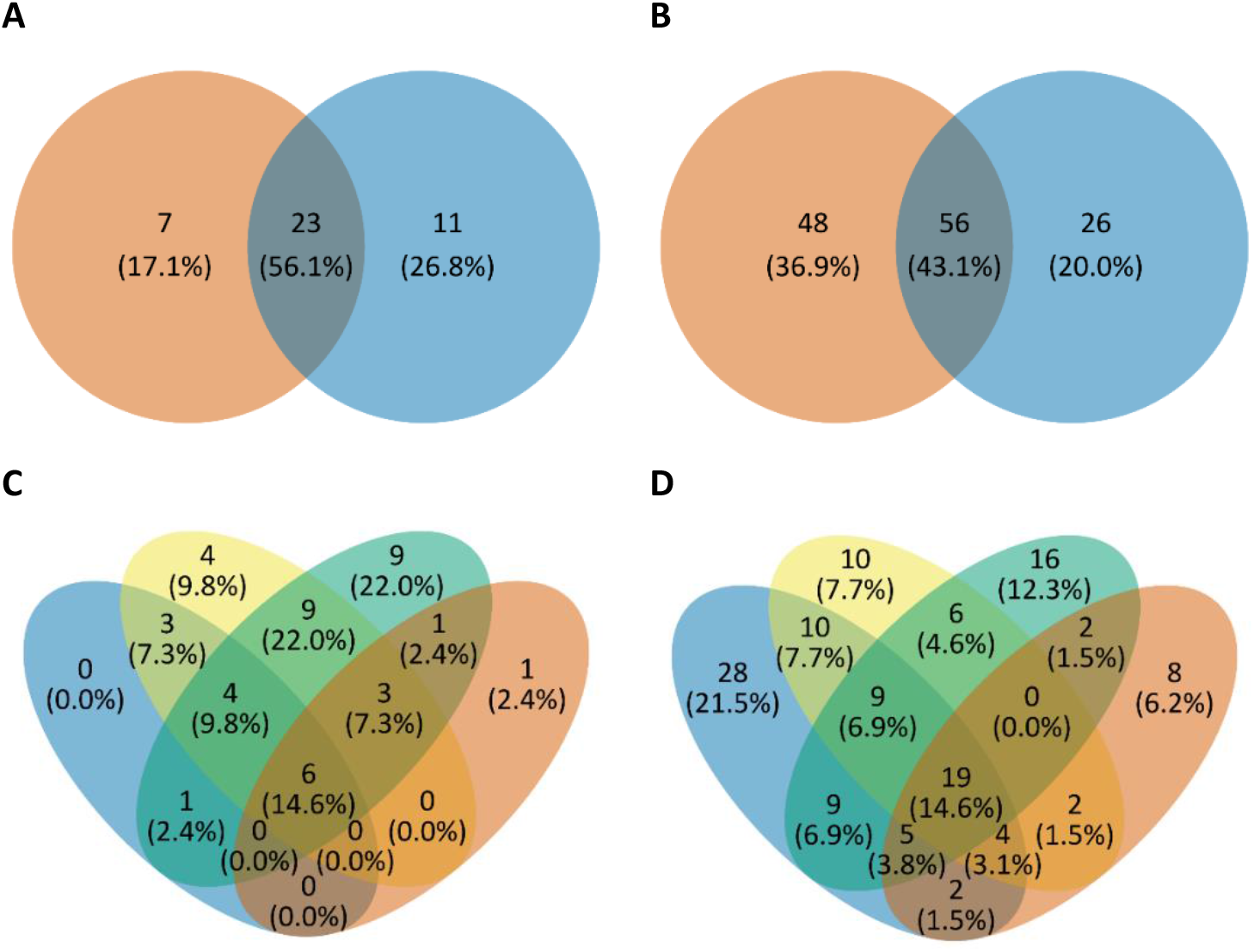
Venn diagrams representing taxa observed within each country using the conventional (A, C) and eDNA (B, D) approaches. A and B represent combined taxa observed per country and C and D represent taxa per habitat per country. Colour representation A and B: orange, Askö; blue, Tvärminne. Colour representation C and D: blue, Askö bare-sediment; yellow, Askö *Fucus;* green, Tvärminne *Fucus;* orange, Tvärminne bare-sediment.

**S7:** Taxa presence per country using the conventional approach.

| Country | AS | TZ |
| --- | --- | --- |
| **ANNELIDA** |  |  |
| *Bylgides sarsi* (Kinberg in Malmgren) | - | Present |
| *Manayunkia aestuarina* (Bourne) | - | Present |
| *Marenzelleria* sp. | Present | Present |
| *Hediste diversicolor* | Present | Present |
| Oligochaeta | Present | Present |
| **ARTHROPODA** |  |  |
| *Asellus aquaticus* (Linnaeus) | - | Present |
| *Balanus* spp. | Present | Present |
| *Chironomus* sp. | Present | Present |
| Copepoda | Present | - |
| Diptera sp | Present | Present |
| *Gammarus* sp | Present | Present |
| *Idotea balthica* (Pallas) | - | Present |
| *Idotea chelipes* (Pallas) | Present | - |
| *Idotea* sp | Present | - |
| *Jaera* sp. | - | Present |
| Ostracoda | Present | Present |
| *Palaemon elegans* (Rathke) | - | Present |
| Plecoptera | - | Present |
| Tanaididae | Present | - |
| Trichoptera | Present | Present |
| Unclassified Insecta | - | Present |
| Unclassified Insecta larvae | - | Present |
| Zygoptera | Present | Present |
| **CHORDATA** |  |  |
| *Gasterosteus aculeatus* (Linnaeus) | Present | Present |
| *Neogobius melanostomus* (Pallas) | - | Present |
| Unclassified fish larvae | Present | - |
| **MOLLUSCA** |  |  |
| *Bithynia* sp. | Present | Present |
| *Cerastoderma edule* (Linnaeus) | Present | Present |
| *Lymnaea stagnalis* (Linnaeus) | Present | Present |
| *Macoma balthica* (Linnaeus) | Present | Present |
| *Mya arenaria* (Linnaeus) | Present | - |
| *Mytilus* spp. complex | Present | Present |
| *Parvicardium* spp. | Present | Present |
| *Peringia ulvae* (Pennant) | Present | Present |
| *Potamopyrgus antipodarum* (J. E. Gray) | - | Present |
| *Radix* spp. | Present | Present |
| *Theodoxus fluviatilis* (Linnaeus) | Present | Present |
| Unclassified gastropod | Present | Present |
| **NEMERTEA** |  |  |
| *Cyanophthalma obscura* (Schultze) | Present | Present |
| **PLATYHELMINTHES** |  |  |
| Platyhelminthes | Present | - |

Abbreviations: AS, Askö; TZ, Tvärminne.

## Notes

### Competing Interest Statement

The authors have declared no competing interest.

## Reference list

Aarnio, K., Mattila, J., 2000. Predation by juvenile *Platichthys flesus* (L.) on shelled prey species in a bare sand and a drift algae habitat., in: Jones, M.B., Azevedo, J.M.N., Neto, A.I., Costa, A.C., Martins, A.M.F. (Eds.), Island, Ocean and Deep-Sea Biology. Developments in Hydrobiology. Springer, Dordrecht. https://doi.org/10.1007/978-94-017-1982-7_32

Anderson, M.J., Gorley, R.N., Clarke, K.R., 2008. PERMANOVA + for FRIMER: Guide to Software and Statistical Methods. PRIMER-E.

Arroyo, N.L., Aarnio, K., Bonsdorff, E., 2006. Drifting algae as a means of re-colonizing defaunated sediments in the Baltic Sea. A short-term microcosm study. Hydrobiologia 554, 83–95. https://doi.org/10.1007/S10750-005-1008-5

Attard, K.M., Rodil, I.F., Berg, P., Norkko, J., Norkko, A., Glud, R.N., 2019. Seasonal metabolism and carbon export potential of a key coastal habitat: The perennial canopy-forming macroalga *Fucus vesiculosus*. Limnol. Oceanogr. 64, 149–164. https://doi.org/10.1002/LNO.11026

Barnes, M.A., Turner, C.R., Jerde, C.L., Renshaw, M.A., Chadderton, W.L., Lodge, D.M., 2014. Environmental conditions influence eDNA persistence in aquatic systems. Environ. Sci. Technol. 48, 1819–1827. https://doi.org/10.1021/ES404734P

Beng, K.C., Corlett, R.T., 2020. Applications of environmental DNA (eDNA) in ecology and conservation: opportunities, challenges and prospects. Biodivers. Conserv. 29, 2089–2121. https://doi.org/10.1007/S10531-020-01980-0

Boström, C., Bonsdorff, E., 1997. Community structure and spatial variation of benthic invertebrates associated with *Zostera marina* (L.) beds in the northern Baltic Sea. J. Sea Res. 37, 153–166. https://doi.org/10.1016/S1385-1101(96)00007-X

Cacabelos, E., Olabarria, C., Incera, M., Troncoso, J.S., 2010. Effects of habitat structure and tidal height on epifaunal assemblages associated with macroalgae. Estuar. Coast. Shelf Sci. 89, 43–52. https://doi.org/10.1016/J.ECSS.2010.05.012

Callahan, B.J., McMurdie, P.J., Rosen, M.J., Han, A.W., Johnson, A.J.A., Holmes, S.P., 2016. DADA2: High-resolution sample inference from Illumina amplicon data. Nat. Methods 13, 581–583. https://doi.org/10.1038/NMETH.3869

El-Khaled, Y.C., Daraghmeh, N., Tilstra, A., Roth, F., Huettel, M., Rossbach, F.I., Casoli, E., Koester, A., Beck, M., Meyer, R., Plewka, J., Schmidt, N., Winkelgrund, L., Merk, B., Wild, C., 2022. Fleshy red algae mats act as temporary reservoirs for sessile invertebrate biodiversity. Commun. Biol. 5, 1–11. https://doi.org/10.1038/S42003-022-03523-5

Elmgren, R., Hill, C., 1997. Ecosystem function at low biodiversity—the Baltic example, in: Ormond, R.F.G., Gage, J.D. (Eds.), Marine Biodiversity: Patterns and Processes. Cambridge University Press, Cambridge, UK, pp. 319–336. https://doi.org/10.1017/CBO9780511752360.015

Engkvist, R., Malm, T., Tobiasson, S., 2000. Density dependent grazing effects of the isopod *Idotea baltica* Pallas on *Fucus vesiculosus* L in the Baltic Sea. Aquat. Ecol. 34, 253–260. https://doi.org/10.1023/A:1009919526259

Everett, R.A., 1994. Macroalgae in marine soft-sediment communities: effects on benthic faunal assemblages. J. Exp. Mar. Bio. Ecol. 175, 253–274. https://doi.org/10.1016/0022-0981(94)90030-2

Foster, N.R., van Dijk, K. jent, Biffin, E., Young, J.M., Thomson, V.A., Gillanders, B.M., Jones, A.R., Waycott, M., 2022. A targeted capture approach to generating reference sequence databases for chloroplast gene regions. Ecol. Evol. 12, e8816. https://doi.org/10.1002/ECE3.8816

Furman, E., Pihlajamäki, M., Välipakka, P., Myrberg, K., 2014. The Baltic Sea–Environment and Ecology. Helsinki, Finland.

Gao, C.H., Yu, G., Cai, P., 2021. ggVennDiagram: An Intuitive, Easy-to-Use, and Highly Customizable R Package to Generate Venn Diagram. Front. Genet. 12. https://doi.org/10.3389/FGENE.2021.706907

Guillou, L., Bachar, D., Audic, S., Bass, D., Berney, C., Bittner, L., Boutte, C., Burgaud, G., De Vargas, C., Decelle, J., Del Campo, J., Dolan, J.R., Dunthorn, M., Edvardsen, B., Holzmann, M., Kooistra, W.H.C.F., Lara, E., Le Bescot, N., Logares, R., Mahé, F., Massana, R., Montresor, M., Morard, R., Not, F., Pawlowski, J., Probert, I., Sauvadet, A.L., Siano, R., Stoeck, T., Vaulot, D., Zimmermann, P., Christen, R., 2013. The Protist Ribosomal Reference database (PR2): a catalog of unicellular eukaryote small sub-unit rRNA sequences with curated taxonomy. Nucleic Acids Res. 41. https://doi.org/10.1093/NAR/GKS1160

Hansen, J.P., Wikström, S.A., Axemar, H., Kautsky, L., 2011. Distribution differences and active habitat choices of invertebrates between macrophytes of different morphological complexity. Aquat. Ecol. 45, 11–22. https://doi.org/10.1007/S10452-010-9319-7

Harper, L.R., Buxton, A.S., Rees, H.C., Bruce, K., Brys, R., Halfmaerten, D., Read, D.S., Watson, H. V., Sayer, C.D., Jones, E.P., Priestley, V., Mächler, E., Múrria, C., Garcés-Pastor, S., Medupin, C., Burgess, K., Benson, G., Boonham, N., Griffiths, R.A., Lawson Handley, L., Hänfling, B., 2019. Prospects and challenges of environmental DNA (eDNA) monitoring in freshwater ponds. Hydrobiologia 826, 25–41. https://doi.org/10.1007/S10750-018-3750-5

HELCOM, 2013. Red List of Baltic Sea underwater biotopes, habitats and biotope complexes, Baltic Sea Environmental Proceedings. Painomies, Finland, Helsinki.

Henseler, C., Nordström, M.C., Törnroos, A., Snickars, M., Pécuchet, L., Lindegren, M., Bonsdorff, E., 2019. Coastal habitats and their importance for the diversity of benthic communities: A species-and trait-based approach. Estuar. Coast. Shelf Sci. 226, 106272. https://doi.org/10.1016/J.ECSS.2019.106272

Huson, D.H., Beier, S., Flade, I., Górska, A., El-Hadidi, M., Mitra, S., Ruscheweyh, H.J., Tappu, R., 2016. MEGAN community edition - Interactive exploration and analysis of large-scale microbiome sequencing data. PLOS Comput. Biol. 12, e1004957. https://doi.org/10.1371/JOURNAL.PCBI.1004957

Huston, M.A., DeAngelis, D.L., 1994. Competition and coexistence: The effects of resource transport and supply rates. Am. Nat. 144, 954–977. https://doi.org/10.1086/285720

Jane, S.F., Wilcox, T.M., Mckelvey, K.S., Young, M.K., Schwartz, M.K., Lowe, W.H., Letcher, B.H., Whiteley, A.R., 2015. Distance, flow and PCR inhibition: eDNA dynamics in two headwater streams. Mol. Ecol. Resour. 15, 216–227. https://doi.org/10.1111/1755-0998.12285

Johannesson, K., Smolarz, K., Grahn, M., André, C., 2011. The future of Baltic Sea populations: Local extinction or evolutionary rescue? Ambio 40, 179–190. https://doi.org/10.1007/S13280-010-0129-X

Johnson, M.T.J., Agrawal, A.A., 2005. Plant genotype and environment interact to shape a diverse arthropod community on evening primrose (*Oenothera biennis*). Ecology 86, 874–885. https://doi.org/10.1890/04-1068

Kangas, P., Autio, H., Hällfors, G., Luther, H., Niemi, Å., Salemaa, H., 1982. A general model of the decline of *Fucus vesiculosus* at Tvärminne, south coast of Finland in 1977–81. Acta Bot. Fenn. 118, 1–27.

Korpinen, S., Honkanen, T., Vesakoski, O., Hemmi, A., Koivikko, R., Loponen, J., Jormalainen, V., 2007. Macroalgal communities face the challenge of changing biotic interactions: Review with focus on the Baltic Sea. Ambio 36, 203–211. https://doi.org/10.1579/0044-7447(2007)36[203:MCFTCO]2.0.CO;2

Kostylev, V.E., Erlandsson, J., Mak, Y.M., Williams, G.A., 2005. The relative importance of habitat complexity and surface area in assessing biodiversity: Fractal application on rocky shores. Ecol. Complex. 2, 272–286. https://doi.org/10.1016/J.ECOCOM.2005.04.002

Kotta, J., Orav, H., 2001. Role of benthic macroalgae in regulating macrozoobenthic assemblages in the Väinameri (north-eastern Baltic Sea). Ann. Zool. Fennici 38, 163–171.

Kovalenko, K.E., Thomaz, S.M., Warfe, D.M., 2012. Habitat complexity: Approaches and future directions. Hydrobiologia 685, 1–17. https://doi.org/10.1007/S10750-011-0974-Z

Kraufvelin, P., Salovius, S., 2004. Animal diversity in Baltic rocky shore macroalgae: Can *Cladophora glomerata* compensate for lost *Fucus vesiculosus?* Estuar. Coast. Shelf Sci. 61, 369–378. https://doi.org/10.1016/J.ECSS.2004.06.006

Kraufvelin, P., Salovius, S., Christie, H., Moy, F.E., Karez, R., Pedersen, M.F., 2006. Eutrophication-induced changes in benthic algae affect the behaviour and fitness of the marine amphipod *Gammarus locusta*. Aquat. Bot. 84, 199–209. https://doi.org/10.1016/J.AQUABOT.2005.08.008

Lauringson, V., Kotta, J., 2006. Influence of the thin drift algal mats on the distribution of macrozoobenthos in Kõiguste Bay, NE Baltic Sea. Hydrobiologia 554, 97–105. https://doi.org/10.1007/S10750-005-1009-4

Levin, S.A., 1992. The problem of pattern and scale in ecology. Ecology 73, 1943–1967.

Levins, R., 1979. Coexistence in a variable environment. Am. Nat. 114, 765–783.

Levy-Booth, D.J., Campbell, R.G., Gulden, R.H., Hart, M.M., Powell, J.R., Klironomos, J.N., Pauls, K.P., Swanton, C.J., Trevors, J.T., Dunfield, K.E., 2007. Cycling of extracellular DNA in the soil environment. Soil Biol. Biochem. 39, 2977–2991. https://doi.org/10.1016/J.SOILBIO.2007.06.020

Lüning, K., 1990. Seaweeds: their environment, biogeography, and ecophysiology. John Wiley & Sons, New York, USA.

Martin, M., 2011. Cutadapt removes adapter sequences from high-throughput sequencing reads. EMBnet.journal 17, 10–12. https://doi.org/10.14806/EJ.17.1.200

Matthäus, W., 2006. The history of investigation of salt water inflows into the Baltic Sea-from the early beginning to recent results. Inst. für Ostseeforschung. https://doi.org/10.12754/MSR-2006-0065

Mosbahi, N., Pezy, J.-P., Dauvin, J.-C., Neifar, L., 2016. Spatial and temporal structures of the macrozoobenthos from the intertidal zone of the Kneiss Islands (central Mediterranean Sea). Open J. Mar. Sci. 06, 223–237. https://doi.org/10.4236/OJMS.2016.62018

Norkko, A., Bonsdorff, E., 1996a. Rapid zoobenthic community responses to accumulations of drifting algae. Mar. Ecol. Prog. Ser. 131, 143–157. https://doi.org/10.3354/MEPS131143

Norkko, A., Bonsdorff, E., 1996b. Population responses of coastal zoobenthos to stress induced by drifting algal mats. Mar. Ecol. Prog. Ser. 140, 141–151. https://doi.org/10.3354/MEPS140141

Norkko, J., Bonsdorff, E., Norkko, A., 2000. Drifting algal mats as an alternative habitat for benthic invertebrates: Species specific responses to a transient resource. J. Exp. Mar. Bio. Ecol. 248, 79–104. https://doi.org/10.1016/S0022-0981(00)00155-6

Ojaveer, H., Jaanus, A., MacKenzie, B.R., Martin, G., Olenin, S., Radziejewska, T., Telesh, I., Zettler, M.L., Zaiko, A., 2010. Status of biodiversity in the Baltic Sea. PLoS One 5, e12467. https://doi.org/10.1371/JOURNAL.PONE.0012467

Oksanen, J., Guillaume Blanchet, F., Friendly, M., Kindt, R., Legendre, P., McGlinn, D., Minchin, P.R., O’Hara, R.B., Simpson, G., Solymos, P., Stevens, M., Szoecs, E., Wagner, H., 2022. vegan: community ecology package. R package version 2.5–7. 2020.

Orav, H., Kotta, J., Martin, G., 2000. Factors affecting the distribution of benthic invertebrates in the phytal zone of the north-eastern Baltic Sea, in: Proceedings of the Estonian Academy of Sciences, Biology Ecology. pp. 253–269.

Pianka, E., 2011. Evolutionary ecology. Harper and Row, New York.

Pietramellara, G., Ascher, J., Borgogni, F., Ceccherini, M.T., Guerri, G., Nannipieri, P., 2009. Extracellular DNA in soil and sediment: Fate and ecological relevance. Biol. Fertil. Soils 45, 219–235. https://doi.org/10.1007/S00374-008-0345-8

Preston, R., Blomster, J., Schagerström, E., Seppä, P., 2022a. Clonality, polyploidy and spatial population structure in Baltic Sea *Fucus vesiculosus*. Ecol. Evol. 12, e9336. https://doi.org/10.1002/ECE3.9336

Preston, R., Rodil, I.F., 2023. Genetic characteristics influence the phenotype of marine macroalga *Fucus vesiculosus* (Phaeophyceae). Ecol. Evol. 13, e9788. https://doi.org/10.1002/ECE3.9788

Preston, R., Rodil, I.F., 2022. Microsatellite genotyping and morphology datasets of Baltic Sea *Fucus vesiculosus*. figshare. Dataset. https://doi.org/10.6084/m9.figshare.19690930.v1

Preston, R., Seppä, P., Schagerström, E., Blomster, J., 2022b. Phylogeographic patterns in attached and free-living marine macroalga *Fucus vesiculosus* (Fucaceae, Phaeophyceae) in the Baltic Sea. Bot. Mar. 65, 419–432. https://doi.org/10.1515/BOT-2022-0016

R Core Team, 2021. R: A language and environment for statistical computing. R Found. Stat. Comput.

Rabalais, N.N., Díaz, R.J., Levin, L.A., Turner, R.E., Gilbert, D., Zhang, J., 2010. Dynamics and distribution of natural and human-caused hypoxia. Biogeosciences 7, 585–619. https://doi.org/10.5194/BG-7-585-2010

Rinne, H., Blanc, J.-F., Salo, T., Nordström, M.C., Salmela, N., Salovius-Laurén, S., 2022. Variation in *Fucus vesiculosus* associated fauna along a eutrophication gradient. Estuar. Coast. Shelf Sci. 107976. https://doi.org/10.1016/J.ECSS.2022.107976

Rognes, T., Flouri, T., Nichols, B., Quince, C., Mahé, F., 2016. VSEARCH: a versatile open source tool for metagenomics. PeerJ 4. https://doi.org/10.7717/PEERJ.2584

Rossbach, F.I., Casoli, E., Beck, M., Wild, C., 2021. Mediterranean red macro algae mats as habitat for high abundances of serpulid polychaetes. Diversity 13, 265. https://doi.org/10.3390/D13060265

Rossbach, F.I., Merk, B., Wild, C., 2022. High diversity and abundance of Foraminifera associated with Mediterranean benthic red algae mats. Diversity 14, 21. https://doi.org/10.3390/D14010021

Roussel, J.M., Paillisson, J.M., Tréguier, A., Petit, E., 2015. The downside of eDNA as a survey tool in water bodies. J. Appl. Ecol. 52, 823–826. https://doi.org/10.1111/1365-2664.12428

Schagerström, E., Forslund, H., Kautsky, L., Pärnoja, M., Kotta, J., 2014. Does thalli complexity and biomass affect the associated flora and fauna of two co-occurring *Fucus* species in the Baltic Sea? Estuar. Coast. Shelf Sci. 149, 187–193. https://doi.org/10.1016/J.ECSS.2014.08.022

Schrader, C., Schielke, A., Ellerbroek, L., Johne, R., 2012. PCR inhibitors – occurrence, properties and removal. J. Appl. Microbiol. 113, 1014–1026. https://doi.org/10.1111/J.1365-2672.2012.05384.X

Stadhouders, R., Pas, S.D., Anber, J., Voermans, J., Mes, T.H.M., Schutten, M., 2010. The effect of primer-template mismatches on the detection and quantification of nucleic acids using the 5’ nuclease assay. J. Mol. Diagnostics 12, 109–117. https://doi.org/10.2353/JMOLDX.2010.090035

Stoeck, T., Bass, D., Nebel, M., Christen, R., Jones, M.D.M., Breiner, H.W., Richards, T.A., 2010. Multiple marker parallel tag environmental DNA sequencing reveals a highly complex eukaryotic community in marine anoxic water. Mol. Ecol. 19, 21–31.

Taberlet, P., Coissac, E., Hajibabaei, M., Rieseberg, L.H., 2012. Environmental DNA. Mol. Ecol. 21, 1789–1793. https://doi.org/10.1111/J.1365-294X.2012.05542.X

Wickham, H., 2016. ggplot2: elegant graphics for data analysis. Springer-Verlag, New York.

Wikström, S.A., Kautsky, L., 2007. Structure and diversity of invertebrate communities in the presence and absence of canopy-forming *Fucus vesiculosus* in the Baltic Sea. Estuar. Coast. Shelf Sci. 72, 168–176. https://doi.org/10.1016/J.ECSS.2006.10.009

Zaiko, A., Pochon, X., Garcia-Vazquez, E., Olenin, S., Wood, S.A., 2018. Advantages and limitations of environmental DNA/RNA tools for marine biosecurity: Management and surveillance of non-indigenous species. Front. Mar. Sci. 5, 322. https://doi.org/10.3389/FMARS.2018.00322

Zettler, M.L., Karlsson, A., Kontula, T., Gruszka, P., Laine, A.O., Herkül, K., Schiele, K.S., Maximov, A., Haldin, J., 2014. Biodiversity gradient in the Baltic Sea: A comprehensive inventory of macrozoobenthos data. Helgol. Mar. Res. 68, 49–57. https://doi.org/10.1007/s10152-013-0368-x

Zhang, Z., Schwartz, S., Wagner, L., Miller, W., 2000. A greedy algorithm for aligning DNA sequences. J. Comput. Biol. 7, 203–214. https://doi.org/10.1089/10665270050081478

Zillén, L., Conley, D.J., Andrén, T., Andrén, E., Björck, S., 2008. Past occurrences of hypoxia in the Baltic Sea and the role of climate variability, environmental change and human impact. Earth-Science Rev. 91, 77–92. https://doi.org/10.1016/J.EARSCIREV.2008.10.001

